# Spatial Single-Cell Mapping of Transcriptional Differences Across Genetic Backgrounds in Mouse Brains

**DOI:** 10.1101/2024.10.08.617260

**Authors:** Zachary Hemminger, Gaby Tam, Haley De Ocampo, Aihui Wang, Thomas Underwood, Fangming Xie, Qiuying Zhao, Dongyuan Song, Jingyi Jessica Li, Hongwei Dong, Roy Wollman

## Abstract

Genetic variation can alter organ structure and, in turn, function. Comparative statistical analysis of organs across genetic backgrounds requires spatial, single-cell, atlas-scale data in replicates, which current technologies do not provide at scale. We introduce **A**tlas-scale **T**ranscriptome **L**ocalization using **A**ggregate **S**ignatures (ATLAS), a scalable tissue mapping method. ATLAS learns transcriptional signatures from scRNAseq data, encodes them *in situ* with tens of thousands of oligonucleotide probes, and decodes them to infer cell types and imputed transcriptomes. We validated ATLAS in the mouse brain by comparing its cell type inferences with direct MERFISH measurements of marker genes and quantitative comparisons to four other technologies. Using ATLAS, we mapped the central brains of five male and five female C57BL/6J (B6) mice and five male BTBR T+ tf/J (BTBR) mice, an idiopathic model of autism, collectively profiling 40 million cells across over 400 coronal sections. Our analysis revealed over 40 significant differences in cell type distributions and identified 16 regional composition changes across male-female and B6-BTBR comparisons. ATLAS thus enables systematic comparative studies, facilitating organ-level structure-function analysis of disease models.

## Main

The quest to link genotype to phenotype is a cornerstone of biology ^1^. Advances in molecular mapping now allow for the measurement of complex phenotypes, such as the spatial organization of cellular transcriptional states in organs ^2^. Mapping multiple genetic backgrounds will deepen our understanding of how genetics shape organ structure, especially for neurodevelopmental conditions like autism, which have strong genetic underpinnings ^3^. However, atlas creation remains limited to single genetic backgrounds or individual animals due to cost and resource constraints ^4,5^, restricting our understanding of how genetic variation impacts organ architecture. Just as genome sequencing became more feasible with the availability of reference genomes, the availability of spatial reference atlases now enables the development of higher throughput approaches for systematic exploration of the relationship between genetics, anatomy, and physiology.

Spatial transcriptomics is the core technology for organ mapping ^2,6^, but its ability to capture all RNA molecules is inherently limited. The ease of mapping nucleic acids and the wealth of information the transcriptome provides about cellular phenotypes ^7^ have made spatial transcriptomics the leading method for organ-wide studies. However, given the enormous number of mRNA molecules, ranging from 10¹⁶ in large human organs like the liver or brain to 10¹³ in smaller organs like the mouse brain, counting every mRNA molecule is impractical, even when only a subset of an organ is mapped via sectioning. Fortunately, capturing all mRNA molecules is unnecessary. The transcriptome is highly redundant, and most spatial transcriptomic technologies capture less than 1% of total mRNA (#measured / #total transcripts in the transcriptome), relying on computational methods to integrate this data with scRNAseq to build comprehensive atlases. These integration methods have evolved substantially over the past decade, from simple landmark gene approaches ^8,9^ to advanced computational techniques that harmonize scRNAseq and spatial data ^10^. Current mouse brain atlases, built with spatial barcoding ^11^, in situ sequencing ^12^, and hybridization methods ^4,5^, rely on these techniques to create detailed maps.

Given that transcript-level spatial transcriptomics data serves as signatures for data integration, we explored whether more efficient methods could yield more informative signatures. Single-cell transcriptomics tools often reduce the dimensionality of sparse, high-dimensional transcriptomes using techniques like principal component analysis (PCA) ^13^ or non-negative matrix factorization (NMF) ^14^, demonstrating that these reduced-dimensional approximations of transcriptional states serve as powerful signatures for scRNAseq data integration. Currently, such signatures are accessible only in silico, requiring laborious RNA counting. Could these transcriptional signatures be measured directly, bypassing the need for individual gene expression measurements? Previous work using aggregate measurements, where multiple genes are measured together, have demonstrated that transcriptional signatures representing aggregation of expression of multiple genes could be decoded through compressed sensing ^15^ or unsupervised learning to provide key information on tissue ^16^. We realized that this concept can be further extended, and with the appropriate design of an oligonucleotide probe pool, aggregate measurements could directly encode lower-dimensional cellular transcriptional signatures *in situ*. This approach enables the direct measurement of transcriptional states without counting individual RNA molecules. It allows us to measure transcriptional signatures at the cell level, making ATLAS suitable for large-scale measurements and facilitating comprehensive organ mapping.

Here, we introduce ATLAS, a scalable tissue mapping technology that combines *in situ* encoding, based on existing reference atlas data (Figure 1), with in silico decoding to generate cell type maps and imputed transcriptomes (Figure 2). We validated ATLAS by directly measuring marker genes in ATLAS samples and conducting quantitative comparisons with reference atlas data (Figure 2). To demonstrate its scalability, we performed comparative studies of sex dimorphism in B6 mice and a comparison between male B6 and BTBR animals, an autism model. We mapped the central brain region (Common Coordinate Framework (CCF) 4.5-9.5 mm) using 21–49 coronal sections per brain. ATLAS produced detailed cell type maps and imputed transcriptomes for over 40 million cells from 405 sections across 15 animals. Using these data, we reconstructed 3D cell type distributions across the three conditions (Figure 3). Two complementary comparative analyses were conducted: first, identifying cell types with statistically significant differences in spatial distribution—two in the male-female comparison and 41 in the B6-BTBR comparison (Figure 4); and second, identifying differences in overall cell type composition, revealing two regions with differences in male-female comparisons and 14 regions in B6-BTBR comparisons (Figure 5). These results demonstrate the power of ATLAS in revealing how genetic variation influences organ architecture.

### *In situ* encoding of cellular transcriptional signatures

ATLAS expands the capabilities of encoding hybridization from binary codes to linear projection. Encoding hybridizations like MERFISH ^17^ and seqFISH ^18^ rely on binary bipartite codebook matrices, where each matrix element bij represents the mapping between gene *i* and readout *j*, achieved using pools of bivalent DNA oligos that bind both the target RNA and the readout. To extend beyond binary mappings, we designed oligo pools to represent continuous weights wij, where each weight corresponds to the number of probes targeting the RNA of gene *i* for a given readout *j*, allowing hybridization to perform a matrix multiplication of a nonnegative weight matrix with a cellular transcriptome vector. This enables the projection of cellular mRNA into a low-dimensional space *in situ* (Figure 1b). We implemented this using DPNMF, a variant of non-negative matrix factorization ^19,20^, which reconstructs gene expression data while separating predefined labels, producing sparse, low-dimensional factorizations. Using BICCN single-cell transcriptional data and cell type labels ^21^, we derived a DPNMF projection matrix that reduces 6,133 genes into an 18-dimensional transcriptional signature (Extended Figure 1). The choice of 18 dimensions was supported by simulation studies (Extended Figure 2). The matrix was scaled and digitized to match the number of encoding probes binding sites available. DPNMF’s sparsity made this feasible with a total of 28,009 probes in the final oligo pool, enabling efficient *in situ* implementation of a dimensionality reduction operation.

**Figure 1.**
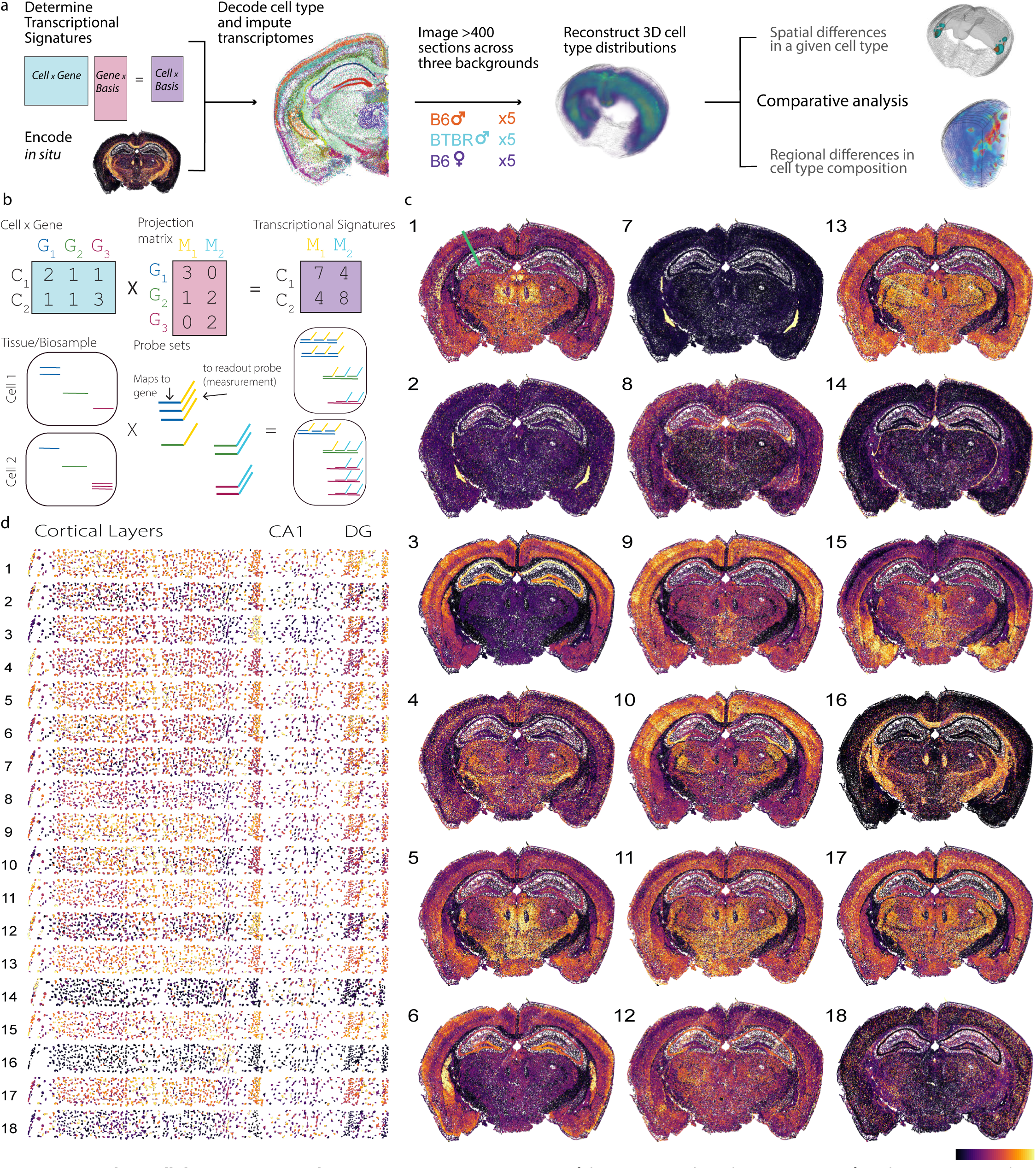
Encoding cellular transcriptional state using ATLAS. (a) Overview of the ATLAS pipeline. (b) Comparison of *in silico* (top row) and *in situ* (bottom row) encoding. The projection matrix, derived from scRNAseq data, is implemented *in situ* by matching the weights of w_ij_ to the number of encoding probes in the oligonucleotide pool. Hybridization, followed by total cell intensity measurements, effectively performs non-negative matrix multiplication. (c) Encoded transcriptional signatures from a single coronal mouse brain section. The green line in (c-1) indicates the region shown at higher magnification in (d), extending from the cortical layer to the dentate gyrus (DG).The colorbar is shown on a logarithmic scale and is independent for each panel, with colors representing relative intensity differences specific to each panel.

To ensure that *in situ* measurements accurately reflected the designed linear projections, we optimized the protocol to minimize non-specific binding and eliminate non-specific signals. Building on the MERFISH protocol ^22^, we made several adjustments (see Methods). Key changes included: (1) Clearing twice, before and after encoder probe hybridization; (2) RNA encoding probes with higher melting temperatures than DNA, allowing them to stay bound during high-temperature clearing and enable more stringent hybridization with DNA readout probes (Extended Figure 3a,b); (3) Enhanced mRNA anchoring to polyacrylamide gels using melpha-X ^23^, ensuring RNA retention during clearing (Extended Figure 3c); and (4) Acquisition of background images before each hybridization round for accurate subtraction of residual fluorescence (Extended Figure 3d). These optimizations enabled us to reliably use total integrated fluorescent intensity to measure cellular transcriptional signatures.

We then stained a brain section with DPNMF probes and measured the projection of cellular transcriptional states (Figure 1c). The resulting data revealed distinct transcriptional states across different cells, which produced brain regions with clear spatial signatures. Some projected components displayed multimodal distributions, while others exhibited more continuous patterns, indicating multiple distinct subpopulations (Extended Figure 4a) providing an insight into DPNMF encoding. Specific regions, such as the cortical layers, demonstrated clear combinatorial encoding of transcriptional states (Figure 1d). These results show that our encoding approach captures rich and diverse information about cellular transcriptional states.

### Decoding and validation of ATLAS signatures

The aggregate signatures represent a dimensionality-reduced version of cellular transcriptional states, making them suitable for direct use in unsupervised clustering. Using standard Leiden clustering on a correlation-based KNN graph of cells (see Methods), we identified distinct clusters that qualitatively align with known brain regions, supporting the notion that ATLAS signatures are highly informative and provide insights into underlying cellular transcriptional states (Extended Figure 4b). This demonstrates the strength of using dimensionality-reduced representations for classifying cells, especially in cases where no agreed-upon cell type taxonomy exists. However, continuously generating new cell type taxonomies with each brain mapping effort is impractical and adds confusion within the research community, as it necessitates mapping across different nomenclatures ^24^. Furthermore, the brain is a unique organ with a very large number of identified cell types, many of which are transcriptionally similar and are primarily distinguished by their spatial location. To address these challenges within the ATLAS framework, we developed a complementary supervised approach that aligns ATLAS’s decoding with existing reference nomenclature, reducing confusion across studies.

To decode ATLAS signatures into predefined cell types, we developed a Bayesian recursive harmonization and classification approach called SCALE (Single Cell Alignment Leveraging Existing data, Extended Figure 5a, see Methods for details). The Bayesian framework was chosen for two key reasons: (1) it maximizes the use of reference atlas information, consistent with the principles of ATLAS, and (2) it leverages prior knowledge of cell type distributions based on anatomical location in the brain. We generated spatial probability maps for each cell type subclass and incorporated supervised learning based on reference scRNAseq data to infer the posterior probability of a transcriptional signature belonging to a specific cell type. The process involved projecting scRNAseq data into a lower-dimensional space and harmonizing it with ATLAS data through a recursive classification and correction approach. Following the strategy used to harmonize reference atlas FISH with scRNAseq data ^4^, we stepped through the cell type dendrogram, applying simple linear corrections after each classification step (Extended Figure 5a,c). This recursive descent brought the ATLAS data into the projected scRNAseq reference space, making imputation a straightforward process using a KNN approach. The final cell type calls qualitatively matched expected data (Figure 2b). We observed batch-to-batch variation in absolute ATLAS signal intensity, whose source is not fully understood, but experimentally measured background correction and median-centering of each latent dimension reduced this variation and improved decoding consistency (Extended Data Fig. 11). We have also shown that the method’s reliance on a spatially-based prior is robust to localization errors of up to hundreds of micrometers (Extended Figure 12).

**Figure 2.**
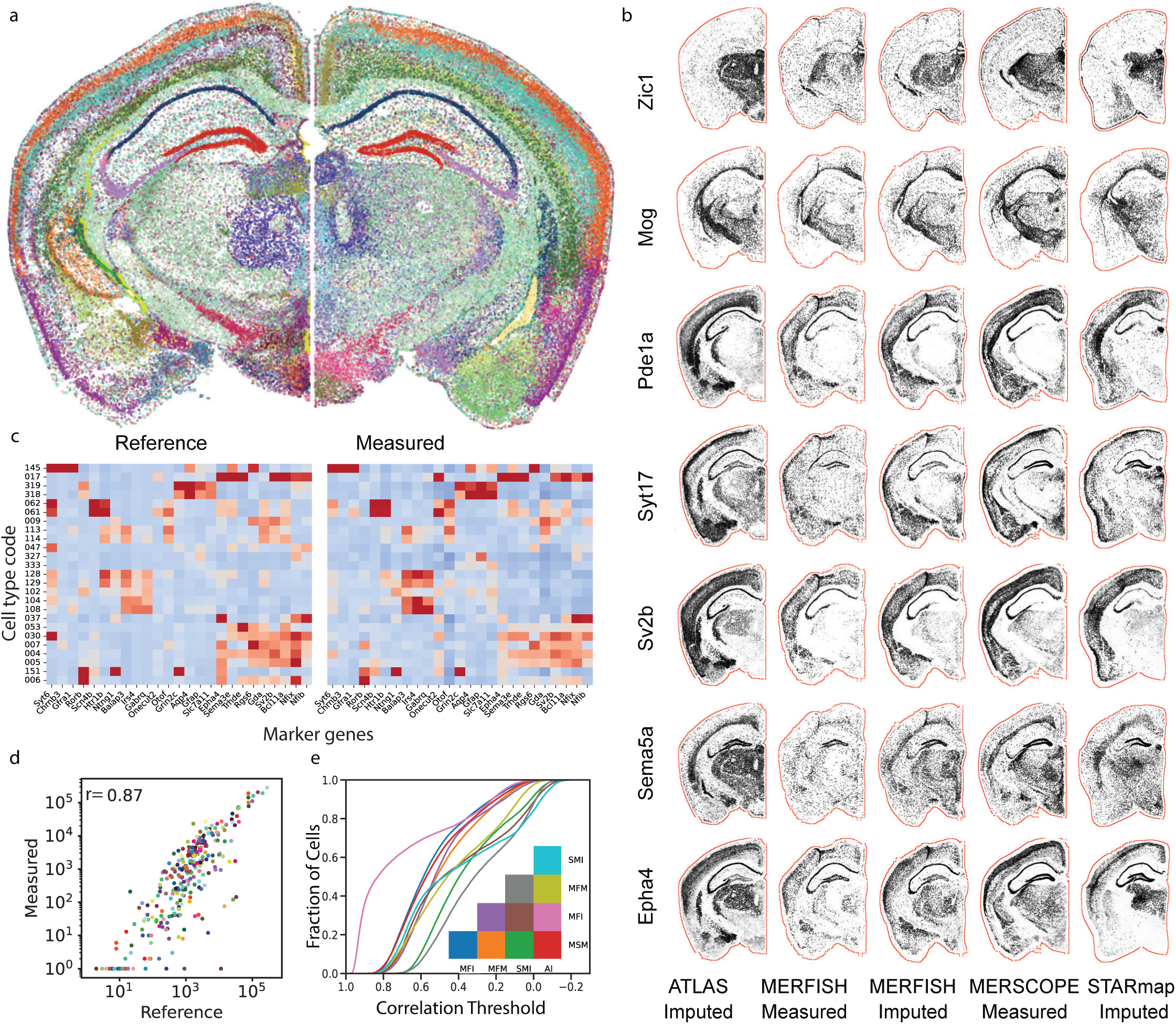
Decoding of ATLAS transcriptional signatures. (a) Comparison between cell types identified using MERFISH gene expression signatures in the reference atlas (left) and cell types inferred from ATLAS transcriptional signatures (right). Cell are colored based on the reference atlas subclass color scheme (b) Representative gene expression plots for seven genes measured across five platforms on matched sections. (c) Comparison of the 25 most abundant cell types, showing reference average expression of marker genes and a validation dataset where cell types were inferred from low-magnification ATLAS imaging, with gene expression measured using MERFISH on the same sample. (d) Total abundance of cell types in the central brain, comparing reference data to ATLAS-inferred data, with the Pearson correlation coefficient (r = 0.87) shown. (e) Quantification of agreement between technologies: for each pair of technologies (color-coded), cumulative probability distributions of gene expression correlation (across 336 genes) are plotted for synthetically paired cells between the technologies (see Methods). Method accronym: SMI-STARmap imputed (Shi et al 2023), MFM-MERFISH measured (Zhang et al 2023), MFI-MERFISH imputed (Zhang et al 2023), MSM-MERSCOPE measured (Yao et al 2023), AI-ATLAS imputed (this study).

To validate ATLAS, we employed two complementary approaches: direct measurement of marker genes and quantitative comparison to existing reference atlases. Marker genes were measured following a modified MERFISH protocol, where cellular mRNAs were anchored to a polyacrylamide gel (see Methods). After imaging, the MERFISH probes were removed using denaturing conditions, and the same sample was hybridized with ATLAS RNA encoding probes. Because the tissue mRNAs were covalently anchored to the gel, we experienced minimal RNA loss, allowing repeated imaging with both protocols. Using this procedure, we measured the expression of 170 marker genes, a subset of the original set used in the reference atlas ^4^, for cells in a single coronal section hemisphere, and obtained their cell type information using ATLAS encoding/decoding. We found remarkable agreement between the reference and ATLAS-decoded marker genes for all cell types, qualitatively validating ATLAS decoding (Figure 2c). Initial quantification showed a strong correlation in cell type abundance between ATLAS and reference data (Figure 2d Pearson correlation of 0.87, comparable to value from equivalent comparison between MERFISH and MERSCOPE data of 0.91, Extended Figure 5e). However, this does not account for spatial positioning. To quantitatively assess spatial predictions, we compared ATLAS to multiple reference datasets, as none represent a definitive biological ‘ground truth.’ We developed a statistical pairing procedure using a greedy algorithm to optimize correlation scores and spatial proximity (see Methods), generating a distribution of correlation values for each cell pair across 336 shared genes (Figure 2e). ATLAS imputation showed comparable agreement with MERSCOPE and MERFISH atlases, similar to the agreement between these two independently measured datasets. STARmap, which uses *in situ* sequencing and a different scRNAseq dataset for imputation, had the lowest agreement with other methods. Notably, the strongest agreement was observed between ATLAS and MERFISH imputation, demonstrating that 18-dimensional cellular-level transcriptional signatures can be as effective as 1100-dimensional gene-level signatures for imputation purposes. Simulation-based perturbations of segmentation masks indicated that ATLAS cell-type calls were robust to the segmentation errors tested here (Extended Data Fig. 6). Additional validation using gene expression averages for different CCF regions is shown in Extended Figure 7. These validation analyses confirm that ATLAS provides accurate cell type inference and gene expression imputation.

### Scaling acquisition to map 15 mouse brains

Encouraged by the high quality of the ATLAS signatures, we expanded the scale of data collection. Iterative FISH approaches rely on high-magnification imaging and a closed chamber design, imaging one coverslip at a time. To fully leverage ATLAS’ capabilities for large-area imaging with low magnification, we redesigned the fluidics system to use an open chamber design (Extended Figure 8). This allowed for multi-well imaging, where three wells were hybridized while the other three were imaged, enabling continuous imaging and minimizing the impact of hybridization time. We also used large 40 mm coverslips in each of the six wells, accommodating four coronal brain sections per coverslip. A single 3-day imaging run thus captures data from 24 brain sections. Data were collected from 15 brains across three conditions (B6 female, B6 male, BTBR male). Initially, each brain was sectioned into 24 serial sections, spaced 200 µm apart, covering the anterior–posterior coordinate in the Allen Common Coordinate Framework (CCF x) from 4.5 to 9.5 mm. The 20 µm-thick sections were taken in triplicate, allowing us to replace sections that did not meet quality standards (see Methods). For instance, brain WTM05 required three imaging runs due to section failures in the first two, ultimately resulting in 49 sections for this brain. After quality control (see Methods), the dataset comprised over 40 million cells from over 400 sections (Figure 3a, Extended Figure 9).

**Figure 3.**
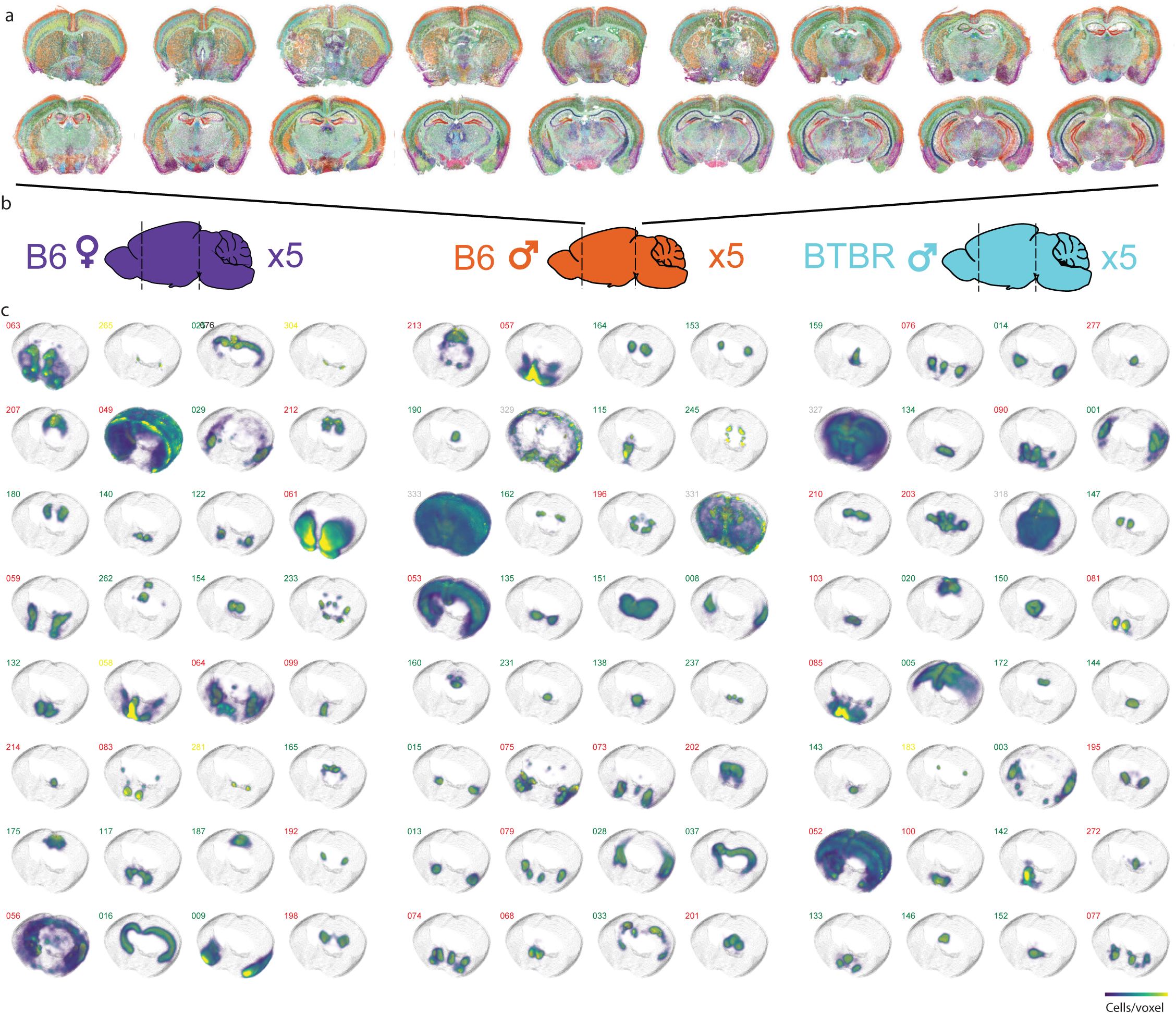
Inference of 3D cell type distributions across B6-female, B6-male, and BTBR-male conditions. (a) Representative 18 serial sections from a single animal, with cell types color-coded based on the reference atlas subclass color scheme. (b) The dataset includes brains from 15 animals, sectioned from CCFx 4.5 to CCFx 9.5, with an average of 27 sections per animal (range: 21–49). (c) Spatial distributions of all 334 cell subclass types were inferred for each condition. Shown are 32 representative cell types per condition. Cell type codes are shown above each distribution color-coded as follows: red (GABA neurons), green (Glutamatergic neurons), yellow (other neurons), and gray (non-neuronal cells). For visualization, different cell types are displayed for each condition, although all 334 types were inferred for all conditions.

This large dataset was then used to construct 3D spatial distribution maps for all 334 cell types at the subclass level across the three mapped conditions (Figure 3c, see Methods for details). In constructing these surfaces, we assumed left-right brain symmetry and pooled information from both hemispheres, yielding ten hemispheric observations from five animals per condition. Pooling information across hemispheres helped address missing values, primarily caused by sectioning artifacts such as tears, folds, etc. Focusing on subclass granularity, we created volumetric estimates of the spatial abundance for each cell type using 100×100×300 µm³ voxels. Encouragingly, the volumetric estimates show high reproducibility between our biological replicas within condition (Extended Figure 10). All statistical comparisons between conditions employed a permutation strategy, where animal labels were permuted, and hundreds of thousands of volumetric cell type spatial abundances were inferred under the null hypothesis that there are no differences between genetic backgrounds.

### Differences in spatial distribution at the cell type level

Building on the accuracy (Figure 2) and scale (Figure 3) of our cell type volumetric maps, we conducted a comparative analysis to identify differences in spatial distributions across B6 male vs. female and B6 male vs. BTBR male. We focused on the spatial distribution of each of the 334 subclasses across the two conditions (see Methods). To ensure that our analysis captured spatially explicit differences, we first calculated for each voxel, the difference (residual) between the two conditions. We then quantified the overall spread of residual distribution using entropy measure. This approach quantifies the divergence between spatial distributions while maintaining spatial context, where lower entropy indicates greater similarity and higher entropy reflects more pronounced differences. Statistical significance was assessed by comparing the observed entropy values to a null distribution generated through permutation. Permutation was performed on the animal label, and the resulting p-values were adjusted for multiple hypothesis testing using the false discovery rate (FDR).

The male-female comparison identified significant spatial differences in two distinct GABAergic neuron populations, providing cellular resolution to the “social behavior network,” a conserved set of interconnected brain regions that includes the amygdala and hypothalamus ^25^ (Figure 4a, Supplementary Table 6). The first cell type, **‘**076 MEA-BST Lhx6 Nfib Gaba,**’** is enriched in the medial amygdala (MEA) and the bed nucleus of the stria terminalis (BST). These core nodes process socio-sexual cues to regulate innate behaviors. The expression of key developmental transcription factors validates this population’s identity: *Lhx6* is a definitive marker for neurons forming a crucial reproductive pathway from the amygdala to the hypothalamus ^26^ while *Nfib* is known to be essential for proper neuronal migration and axonal projection during brain development ^27^. The second population, ‘050 Lamp5 Lhx6 Gaba,’ represents a novel finding localized to the preoptic area (POA), a quintessential sexually dimorphic region that orchestrates male-typical behaviors ^28^. The expression of *Lamp5* is notable, as it canonically marks cortical interneurons that modulate synaptic transmission ^29^, suggesting a potential modulatory mechanism that has been previously unappreciated in the context of sexual dimorphism. Analyzing the magnitude and spatial distribution of these differences (Figure 4c), we found that they are localized to the hypothalamus and affect up to ∼1% of neurons in regions such as the medial amygdala (MEA), indicating a modest but regionally specific impact.

**Figure 4.**
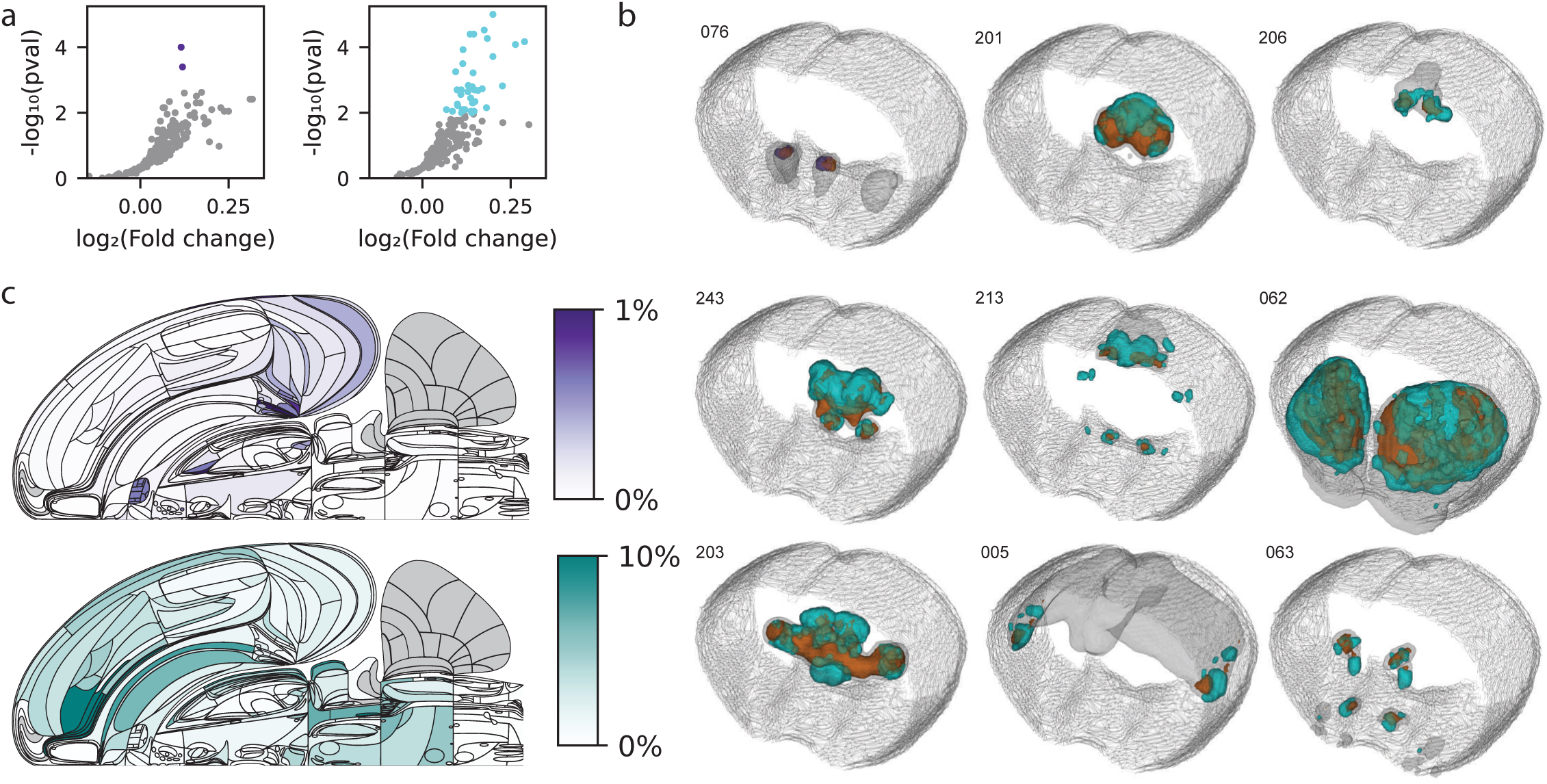
Spatial differences in cell type distribution across male-female and B6-BTBR comparisons. **(a)** Volcano plots illustrating effect size versus significance from the statistical permutation test (see Methods) for all 334 cell types at the subclass level. The comparison between B6 males and B6 females (left) identified two cell types with significant spatial distribution differences, while the B6-BTBR comparison (right) identified 41 cell types. **(b)** Nine representative cell types with significant spatial distribution differences. The overall volume occupied by each cell type across conditions is shown in gray, with areas of increased levels in B6 males, B6 females, or BTBR animals highlighted in orange, purple, and cyan, respectively. Shown (left to right, top to bottom) are: 076_MEA-BST_Lhx6_Nfib_Gaba, 201_PAG-RN_Nkx2-2_Otx1_Gaba, 206_SCm-PAG_Cdh23_Gaba, 243_PGRN-PARN-MDRN_Hoxb5_Glut, 213_SCsg_Gabrr2_Gaba, 062_STR_D2_Gaba, 203_LGv-SPFp-SPFm_Nkx2-2_Tcf7l2_Gaba, 005_L5_IT_CTX_Glut, and 063_STR_D1_Sema5a_Gaba. Colors show the dominant condition (orange for B6 male, purple for B6 female, cyan for BTBR) in each voxel. **(c)** Flatmap representation quantifying the number of cell type changes in each brain region, shown as a weighted fraction of the total number of cell types in the region, adjusted by cell type abundance. Colorbar shows the fraction of total differences over total cells for each brain region summed over all cell types. The top map shows the male-female comparison, while the bottom map shows the B6-BTBR comparison.

The B6-BTBR comparison revealed 41 significant differences in cell type distributions, affecting a larger number of brain regions and showing greater magnitude than the male-female comparison (Figure 4a,b). These differences included both inhibitory and excitatory neuron subclasses. Notably, changes were enriched in the Agranular Insular cortex (AIv and AId), with overall differences in neuronal populations exceeding 10% in these regions. These findings are consistent with previous research that showed that the insular cortex is a critical site of disparity in BTBR mice ^30,31^, and impairment of balance between excitatory and inhibitory neurons in this area underlie the deficits in multisensory integration and the behavioral phenotypes characteristic of the BTBR model of autism spectrum disorder. Another region showing over 10% change was the dorsal raphe nucleus (DR), the brain’s primary source of serotonin. This cellular-level finding provides a critical link to known serotonergic system dysregulation in BTBR mice, which includes altered receptor signaling and a compensatory increase in the number of serotonin-producing neurons within the DR ^32^. These changes likely contribute to the social and anxiety-related behavioral phenotypes characteristic of this model.

These findings underscore the strengths of ATLAS in comparative spatial distribution analysis, revealing both well-characterized and previously unexplored differences in cell type spatial distributions. The high granularity of transcriptionally defined neuron types provided more detailed insights than prior histopathological studies, which reported only modest differences between B6 and BTBR brains ^33^. However, using high-granularity cell types as a comparison unit poses challenges: (1) molecular changes within a cell type may not alter its transcriptional state sufficiently to change ‘type identity,’ and (2) localized shifts in cell type distribution may be difficult to detect when most of the cell type remains spatially consistent. While the first challenge is intrinsic to ATLAS and requires further data, the second could be addressed through complementary analysis that compares spatial distributions on a voxel level rather than by cell type.

### Regional differences in cell type composition

The composition of cell types across brain regions plays a key role in shaping brain function. To assess local and regional differences in cell type composition for the male-female and B6-BTBR comparisons, we calculated the voxel-wise correlation distance of cell type composition vectors between the two conditions. These correlation distances were compared to a null distribution generated by permuting animal labels, resulting in a 3D probabilistic map that highlights spatial locations where differences between conditions are unlikely to arise by chance (Figure 5). Repeating the same compositional analysis after decoding all samples without spatial priors produced highly concordant thresholded p-value maps for both the B6 male versus BTBR male and B6 male versus B6 female comparisons, indicating that the main regional composition results are not driven by the spatial prior (Extended Data Fig. 16).

**Figure 5.**
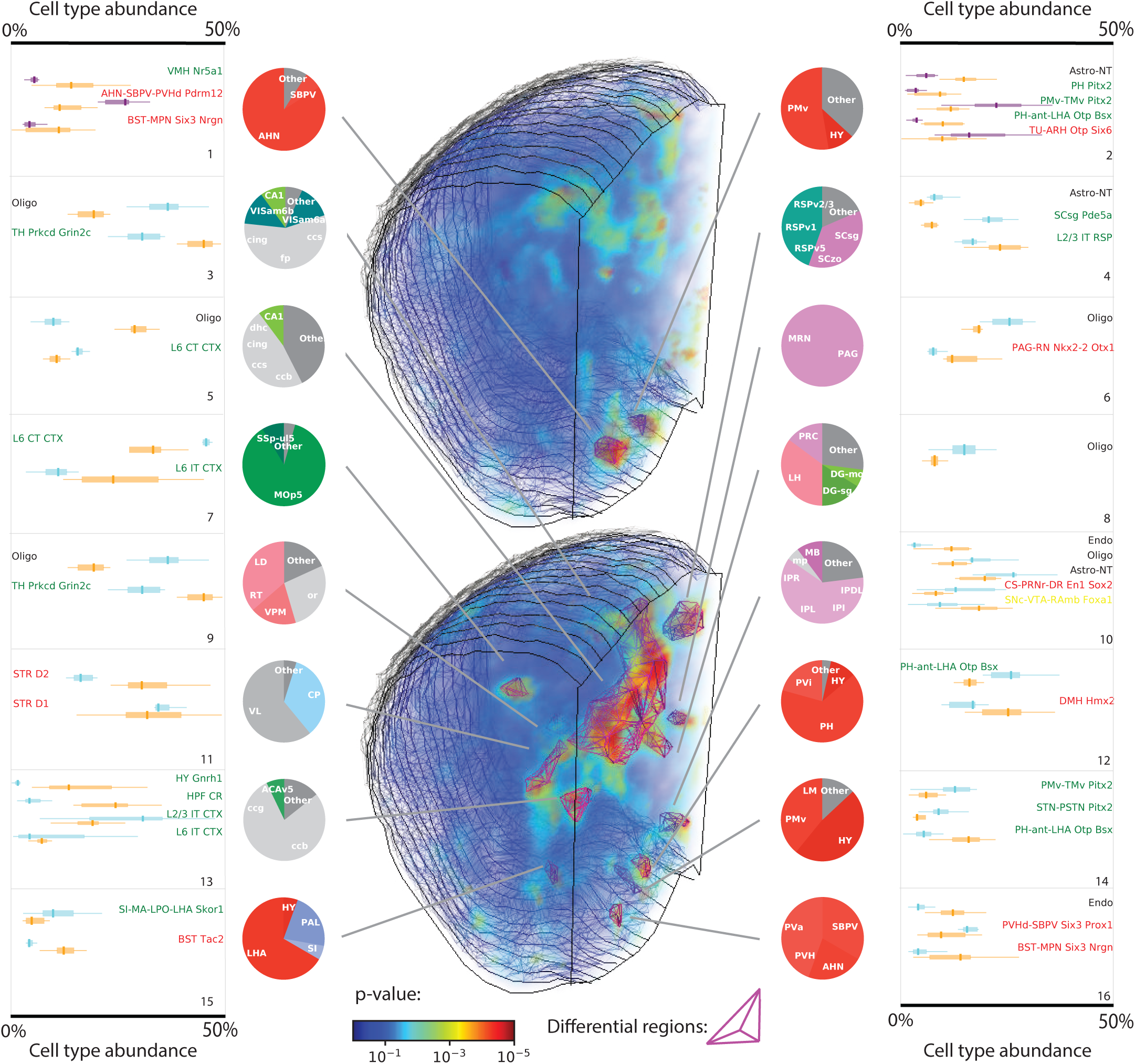
Spatial differences in regional composition across male-female and B6-BTBR comparisons. Volumetric maps show p-values for each spatial position, indicating the likelihood that cell type composition in B6 females (top) or BTBR males (bottom) differs from B6 males, based on permutation testing. Differential regions were defined as contiguous areas larger than 0.015 mm³ with voxel p-values ≤ 0.0002. Two differential regions were identified in the B6 male versus B6 female comparison, while 14 regions were identified in the B6 versus BTBR comparison. For each region, pie charts show the magnitude of overlap between each differential region and known brain areas; acronyms and color codes follow the CCF atlas. Outer panels of each pie chart display cell types in each differential region with >5% abundance difference. Boxplot center line shows the median; box limits show the upper and lower quartiles; whiskers show 1.5x the interquartile range. Colors represent genetic background: orange, B6 male; purple, B6 female; cyan, BTBR male.

We identified two adjacent regions within the hypothalamus that exhibited significant sexual dimorphism. Region 1 overlaps with the anterior hypothalamic nucleus (AHN), while Region 2 primarily overlaps with the ventral premammillary nucleus (PMv). Both regions are known to be sexually dimorphic ^34^, though they have been studied to different extents. Interestingly, the AHN was one of the first sexually dimorphic regions identified in rats in 1977 ^35^ and was initially believed to be similar across mammals. However, follow-up studies showed that, unlike in rats, there are no morphological differences in the AHN of mice ^36^. More recent molecular studies have re-established its sexually dimorphic nature ^37^. Our findings of specific neuronal subclass differences help further refine the molecular resolution of its dimorphic nature and provide a cellular basis for its role in sex-specific behaviors like aggression and parental care. Region 2, the PMv, is a well-established sexually dimorphic region, with roles in maternal aggression ^38^, reproductive control ^39^, male social behavior ^40^, and intermale aggression ^41^. Previous studies identified sexually dimorphic neuron populations within the PMv (e.g., PMv-DAT ^42^, PMv-PACAP ^43^), complementing our systematic identification of subclass-level abundance changes. Interestingly, although we identified several neurons associated with the ventromedial hypothalamus (VMH), a known sexually dimorphic region located between the AHN and PMv, the VMH itself did not show significant differences (p-value = 0.08). Interpreting such negative results is challenging as it could simply be that we do not have the statistical power to find differences and further analysis using the high throughput capabilities of ATLAS may clarify this. It is also possible that sexual dimorphism in the VMH is driven more by gene expression or chromatin state changes than by cell type abundance, an open question in many sexually dimorphic brain regions ^44^.

Our comparison of the BTBR mouse revealed 14 regions with distinct cell compositions, providing cellular-level details that align with and further specify the model’s known neuropathology. Most prominently, we found a complex, bidirectional oligodendrocyte pathology. A marked decrease in oligodendrocytes in fiber tract regions (Regions 3 & 5) is a direct consequence of the BTBR’s hallmark agenesis of the corpus callosum ^33^. These regions also overlapped with areas with high morphological difference from the CCF reference (Extended Figure 14). Conversely, other areas (Regions 6, 8-10) showed a surprising increase in oligodendrocytes, potentially reflecting dysregulated developmental myelination ^45^. This reorganization extended to gray matter, with alterations in the retrosplenial cortex (Region 4) aligning with its known volume reduction as observed by fMRI studies ^46^. The specific findings of cell type compositional changes provide further molecular details of these underlying morphological changes. Crucially, some changes directly inform core ASD-related hypotheses. In the striatum (Region 11), an apparent switch from D1- to D2-type neurons offers a cellular mechanism for the model’s hallmark repetitive behaviors by altering the balance of motor-controlling pathways ^47^. These findings reinforce previous work on changes in dopaminergic receptor distribution in the dorsal striatum ^48^. Similarly, a shift in the cortex (Region 7) from long-range projecting L6 CT to local L6 IT neurons ^49^ provides direct support for the ‘altered connectivity’ hypothesis of ASD ^50,51^. Overall, our analysis identifies distinct cellular pathologies that offer direct mechanistic explanations for the BTBR model’s most prominent features, including its callosal agenesis, repetitive behaviors, and altered brain connectivity.

These findings demonstrate ATLAS’s capability to perform large-scale comparative compositional analyses across spatial volumes exceeding 400 mm³, with voxel sizes of 0.003 mm³, each containing tens of cells. Many of the regions identified as compositionally different were previously associated with sexual dimorphism or behavioral differences between B6 and BTBR mice, while other regions are novel. To experimentally test the predictions stemming from the compositional analysis, we performed direct measurements of cell type abundance using marker genes. Overall, we performed eight comparisons of cell type and regional differences between male B6 and BTBR mice (Extended Figure 15). We found that five of these comparisons showed statistically significant abundance differences, validating our findings and the ability of ATLAS to identify differences in cell type abundance between conditions. Further investigation is required to elucidate the functional implications of these newly identified changes in cellular composition across these regions.

The comparative analysis presented here was performed at the cell type level. To verify that ATLAS contains transcriptional information beyond the cell type label, we performed a spatially-varying differentially expressed gene (spDEG) analysis ^52^. This analysis aims to identify genes whose expression exhibits non-uniform spatial patterns (e.g., patchy distributions or smooth gradients) that cannot be explained by the cell type label alone. Focusing on genes shared with MERSCOPE, a limited analysis of a single coronal section identified numerous cell-type-specific spDEGs, most of which were also detected in a matched MERSCOPE section (Extended Data Fig. 13). However, a targeted comparison to 30 previously reported BTBR-associated genes ^53,54^ showed that BTBR–WT expression differences estimated from ATLAS imputation were not concordant with direct measurements, whereas two direct-measurement datasets were concordant with one another, indicating that WT-reference-based imputation is not a reliable assay for BTBR-associated differential expression (Extended Data Fig. 17).

## Discussion

ATLAS is a scalable approach for mapping cellular transcriptional states by learning transcriptional signatures from reference scRNAseq data, encoding them *in situ* with oligo pools, and decoding them to infer cell types and imputed transcriptomes. By incorporating data integration from the outset, ATLAS optimizes tissue mapping by eliminating the need to measure individual genes, relying instead on transcriptional signatures derived from matrix factorization. We validated ATLAS’s accuracy through direct measurements of marker genes and quantitative comparison with four other brain atlases, demonstrating its performance to be on par with or better than traditional gene expression reconstruction methods. ATLAS’s high throughput, as demonstrated by profiling over 40 million cells across 15 animals, enabled in-depth comparative analyses that revealed significant differences in cell type spatial distributions and regional composition across the brain. These differences range from pinpointing subtle GABAergic subtype differences in sexually dimorphic hypothalamic nuclei to providing cellular-level explanations for the hallmark features of the BTBR autism model. Specifically, ATLAS quantified the bidirectional oligodendrocyte pathology tied to the model’s anatomical defects and, crucially, identified neuronal switches that offer direct mechanistic explanations for behavior. For instance, a shift from D1- to D2-type neurons in the striatum provides a cellular mechanism for the model’s repetitive behaviors by altering motor pathway balance, while a switch from long-range L6 CT to local L6 IT cortical neurons offers a molecular basis for the ‘altered connectivity’ hypothesis of ASD. This capacity for robust, quantitative comparison moves beyond simple mapping, providing a transformative tool for rigorously linking cellular architecture to organ function and dysfunction.

ATLAS introduces a “reference-based assembly” philosophy, designed for an era where foundational reference atlases already exist. Instead of building new atlases from scratch, ATLAS intelligently leverages this existing knowledge at every step. First, its encoding strategy is supervised, using Discriminant Projective Non-Negative Matrix Factorization (DPNMF) on reference scRNA-seq data to design transcriptional signatures explicitly optimized for cell typing. These signatures are then physically measured *in situ* through a unique weighted linear projection, where the number of probes for each gene’s transcript is scaled to match the DPNMF-derived weights. The fidelity of these aggregate measurements is ensured by a suite of experimental optimizations, including enhanced mRNA anchoring with MelphaX and round-specific background subtraction, while a custom high-throughput fluidics system provides the necessary scale. Finally, the decoding process completes this reference-based loop, with the SCALE algorithm using the reference transcriptome and spatial priors to accurately infer cell types and impute gene expression. It is this synergy of reference-guided design, measurement, and decoding that empowers ATLAS to generate comparative spatial atlases at an unprecedented scale.

ATLAS’s scalability arises from its shift from single-molecule to single-cell imaging, resulting in ∼100x faster imaging and ∼100x smaller image datasets, simplifying downstream computational analysis. Importantly, this benefit comes without a reduction in accuracy. However, there are downsides to transitioning from molecule-level to cell-level transcriptional signatures. First, ATLAS lacks subcellular resolution. While this can be addressed by combining ATLAS with MERFISH to capture submicron gene expression data as demonstrated in Figure 2, doing so reintroduces molecule-level measurements, counteracting ATLAS’s key advantages. Second, ATLAS requires high-quality scRNAseq reference data. Although the decreasing cost of scRNAseq and the growing availability of reference datasets mitigate this issue in most cases, challenges remain when mapping nontraditional model organisms or disease conditions with heavily perturbed transcriptional states, where generating new reference data would increase overall costs. Third, ATLAS completely relies on data integration with scRNAseq. While gene-level transcriptional signatures also benefit from data integration, they can still be interpreted directly without it. ATLAS’s deep reliance on data integration is both its greatest advantage—enabling its substantial scalability—and its principal limitation, as it makes the method more indirect. This limitation was evident in the BTBR analysis, where expression values imputed from a WT reference did not reliably recover previously reported BTBR-associated differential expression (Extended Data Fig. 17). This trade-off ensures that there will always be a role for approaches providing direct gene expression measurements.

Several avenues exist for improving ATLAS. First, the reference dataset used to design the encoding scheme was outdated and only included scRNAseq data from the cortex and hippocampus ^21^. The fact that this encoding performed well across other brain regions highlights the robustness of our approach, but updating the reference data would further enhance ATLAS’s accuracy. Second, in this study, we chose to use 18-dimensional transcriptional signatures. This choice represents a tradeoff between accuracy and throughput (Extended Figure 2b), and future applications of ATLAS should adjust this hyperparameter to balance throughput needs and decoding complexity. Third, ATLAS’s throughput could be substantially increased through signal amplification methods, reducing imaging times and potentially eliminating the need to capture background signals, which could cut imaging time in half. However, the amplification must be linear to preserve the accurate representation of cellular transcriptional states, a requirement that does not apply to transcript-level measurements. This limits the use of non-linear amplification methods like RCA ^55^, but both bDNA ^56^ and HCR ^57^ offer promising avenues for linear amplification, potentially boosting throughput by another 10x. Fourth, the polyacrylamide RNA anchoring and extensive clearing used in ATLAS make it compatible with light sheet imaging, which could further increase throughput and enable 3D reconstruction of entire organs. Fifth, in addition to technical improvements, applying ATLAS to organs beyond the brain could offer further advantages; the lower transcriptional diversity typical of many tissues would simplify the encoding challenge and may alleviate the need for spatial priors in decoding. In the brain, we used spatial priors in part because many existing brain cell-type labels are closely tied to anatomical location. However, the regional compositional analysis was largely preserved when decoding was repeated without spatial priors (Extended Data Fig. 16), suggesting that ATLAS can support spatial comparative analyses without requiring the same degree of anatomical prior information. This result is encouraging for extending ATLAS to tissues beyond the brain, where cell-type identity may be less dependent on fine anatomical position. Finally, while this study focused on cell type-related gene expression programs, it is possible that some forms of brain sexual dimorphism and other differences between B6 and BTBR mice were not fully captured in our comparative analysis. This analysis could be extended by leveraging the current imputation or by using ATLAS to encode other gene expression programs involving multiple genes, such as those related to inflammation, cancer, or stress, by incorporating additional aggregate signatures, thereby providing cellular insights beyond cell type identity. These extensions will make ATLAS a robust spatial transcriptomics platform for tissues where an initial reference atlas exists.

Databases like the Protein Data Bank (PDB) have been essential in understanding the structure-function relationship at the molecular level, often through comparisons of wild-type and mutated protein structures. Similarly, ATLAS offers a way to reconstruct anatomical structures at single-cell resolution across conditions with known genetic and functional differences. This paves the way for the creation of an ‘Organ Data Bank,’ analogous to the PDB, that would enable systematic analysis of the relationship between organ structure (anatomy) and function (physiology).

## Supporting information

All Supp Tables

## Data and code availability

All data is available here (https://doi.org/10.5281/zenodo.13851748) and code can be found here (ATLAS https://github.com/wollmanlab/ATLAS_Mouse_Brain/ MERFISH https://github.com/wollmanlab/PySpots Imaging https://github.com/wollmanlab/Scope Fluidics https://github.com/wollmanlab/Fluidics)

## Acknowledgements

This project was supported by NIH grant R01-HG012925.

## Methods

### Encoding Matrix Design

Encoding Matrix was fit using SMART-seq data collected from the mouse whole cortex and hippocampus due to its high capture efficiency per cell ^1^. Gene expression was normalized to a total sum of 100,000. Genes with an average expression below 1 in any cell type were removed to encourage weight assignment to higher expressed genes. Genes that had an average expression above 100 in any cell type were also removed to encourage the use of more genes as highly expressed genes have a strong effect on reconstruction accuracy as well as cell type distances purely due to magnitude. Potential encoding probes were designed for each gene using PaintSHOP ^8^ with a probe length of 30bp. Genes were further filtered if no probes could be designed. Reference dataset was class balanced to the cluster level to ensure that rarer cell types had a meaningful effect on the design. Encoding matrix was fit using a modified version of Discriminant Projective Non Negative Matrix Factorization (DPNMF ^2,3^) which consists of a discriminant aspect that maximizes the variance between cell types while minimizing the variance within a cell type as well as a reconstructive aspect that increases the accuracy of gene reconstruction. A high mu value of 50 was used for the discriminant aspect ensuring that more weight was applied to the discriminant aspect than the reconstruction aspect. The resulting loading matrix was scaled bitwise and integerized to maximize the utilization of the encoding probes that were designed for each gene. This involved clipping the highest weighted genes per bit and scaling so that the highest weighted genes utilized all encoding probes that were possible for those genes. The integerized encoding matrix was then clipped for each gene at the maximum number of probes that could be designed for that gene (i.e. if a gene only had 27 probes but needed a weight of 30 the encoding matrix was clipped to 27). Resulting encoding matrix consists of 6,133 Genes and 28,009 Probes and can be found in Supplementary Table 2.

### Encoding Probe Design

A complex oligo pool of encoding probes was designed for ATLAS consisting of 28,009 encoding probes, targeting 6,133 genes. ATLAS encoding probes were engineered to contain a 30-nucleotide target sequence with specific homology to the mRNA of interest. The targeted sequences for the encoding probes were designed to have a GC content ranging from 45% to 65%, resulting in a melting temperature between 65°C and 72°C. 20-nucleotide readout arms were appended to the ends of the target sequence, and concatenated by two flanking regions designed for primer amplification. Readout sequences and bit assignment can be found in Supplementary Table 5. Two primers, a forward and reverse, were designed for the initial amplification of the oligo pool, with minimal homology, to any encoding probe. The reverse primer was designed with a NheI restriction digest site, used in later steps. An additional forward primer containing a T7 promoter was also designed for subsequent rounds of PCR amplification. Probe sequences can be found in Supplementary Table 3.

For marker gene MERFISH validation probes, an oligo pool of 16,320 encoding probes was designed to target 170 genes. The probe design follows the same structure as the ATLAS probes, with the key difference being that four readout arms were appended to the ends of the target sequences. Additionally, the same forward and reverse primers were incorporated at the ends of the probes. MERFISH codebook which shows the barcode for each gene can be found in Supplementary Table 9. Encoding probe sequences can be found in Supplementary Table 8.

For marker gene RNAFISH validation probes, an oligo pool of 240 encoding probes was designed to target 6 genes with 40 probes for each gene. Probes consisted of a 30mer encoding site consistent with ATLAS probes and four 20mer readout probe binding sites, two on each side of the 30mer. Probes were used directly as IDT oPools and thus did not require primers for amplification. Probe sequences can be found in Supplementary Table 7.

### Encoding Probe Amplification

The template molecules for the ATLAS oligo pool (Twist Biosciences) were amplified in two limited cycle PCR reactions to minimize the formation of nonspecific products. A small-scale PCR reaction was first carried out using 0.4 ng/uL of the initial Twist template, following the manufacturer’s recommended guidelines. KAPA HiFi hot start ready mix (Fisher Scientific, 50-196-5217) was used for PCR amplification, with 0.3 µM of primers. The initial template amplification did not include the forward primer with T7 promoter. The correct product size was validated on an Agilent 2100 bioanalyzer and then on a 15%TBE-Urea polyacrylamide gel (Thermo Fisher Scientific, EC68852BOX) in every step after that. The PCR product was cleaned using a phenol-chloroform extraction and desalted using a 10 kDa centrifugal filter column (Sigma Aldrich, UFC5010). The amplified product was used as the template for the second PCR reaction at 0.04 ng/uL per reaction volume. Product was amplified with KAPA HiFi hot start ready mix and 0.3 µM T7 forward primer, adjusting the melting temperature accordingly. For marker gene validation probes, the second PCR reaction was performed with 0.3 µM T7 reverse primer. Elongation times for the second reaction were also increased to 45 sec, deviating from Twist recommendations of 15 seconds. To enhance probe penetration, the PCR product was digested overnight at 37°C with 1 unit of NheI-HF (New England Biolabs, R3131) per µg of product, reducing the size of the encoding probes from 113 nt to 94 nt.

Digested PCR products were converted to RNA encoding probes using a high yield in vitro transcription (IVT) kit (New England Biolab, E2040S). Reaction concentrations were maintained as per the manufacturer’s recommendations, with the exception of cytidine triphosphate (CTP; Thermo Fisher Scientific, R0451), which was added to the reaction volume at a final concentration of 5 mM. IVT amplification was carried out overnight at 37°C before cleaning and desalting as described above. The final concentration of encoding probes was quantified using a BR RNA Qubit kit (Thermo Fisher Scientific, Q10210) and size verified via gel electrophoresis. RNA encoding probes were aliquoted into 1.5 mL tubes, with 600 μg allocated per experiment. The encoding probes were dried completely using a speedvac and stored at −80°C until further use. The same protocol was followed for the preparation of nonspecific encoding probes.

For marker gene validation probes, IVT products were converted to DNA encoding probes using Reverse Transcriptase (Maxima H Minus Reverse Transcriptase, EP0751). Reaction concentrations were maintained as per the manufacturer’s recommendation with 40 µM forward primer containing a Uracil at the 3’ end. RT was carried out overnight at 53°C before cleaning and desalting as described above. To enhance probe penetration, the RT product was digested overnight at 37°C with 1 unit of USER Enzyme (New England Biolabs, M5505L) per µg of product, reducing the size of the encoding probes from 150 nt to 130 nt. DNA encoding probes were aliquoted into 1.5 mL tubes, with 100 μg allocated per experiment. The encoding probes were dried completely using a speedvac and stored at −80°C until further use.

### Coverslip Functionalization

Coverslips (40 mm, #1.5; Bioptechs, 40-1313-03193) were cleaned by submerging in a 1:1 solution of 37% HCl and methanol with sonication for 30 minutes. They were then rinsed twice with deionized water and once with 100% ethanol, each for 5 minutes, before being dried completely at 70°C. The coverslips were subsequently submerged in a mixture of 0.1% (v/v) triethylamine (Sigma Aldrich, 471283) and 0.2% (v/v) allyltrichlorosilane in chloroform for 30 minutes. After rinsing once in chloroform and twice in 100% ethanol, the coverslips were dried at 70°C. They were then treated with 2% (v/v) (3-Aminopropyl)triethoxysilane (APES, Sigma Aldrich, 440140) in acetone for 10 minutes, followed by two 5 minute rinses in deionized water and once in 100% ethanol. The coverslips were dried once more at 70°C and stored under vacuum until further use. Functionalization treatment is necessary for improved tissue and gel adhesion to coverslips ^4^.

### Animals

Adult BTBR *T*+ *Itpr3tf*/J male and C57Bl6/J male and female mice aged 56 days were used for this study. To reduce the stress of animals due to shipping and handling, mice were maintained for one week upon arrival on a 12 hour:12 hour light/dark cycle with access to food and water before sectioning. All mice were maintained and bred under standard conditions consistent with NIH guidelines and approved by the Chancellor’s Animal Research Committee at the University of California, Los Angeles.

### Sectioning

Whole mouse brains were harvested at 8 weeks of age and perfused promptly in 1x PBS (Thermo Fisher Scientific, 10010049) with 0.1% (v/v) Tween-20 (Sigma Aldrich, P1379) and 3 mg/mL Poly(vinylsulfonic acid, sodium salt) solution (PVSA, Sigma Aldrich, 278424) (1x PBSTw). Samples were embedded in optimal cutting temperature (OCT, Fisher Scientific, 23-730-571) compound immediately and flash frozen in liquid nitrogen before being stored at −80°C. The day before sectioning, samples were mounted onto a cryostat specimen disk and stored at −20°C allowing the sample to equilibrate to sectioning temperatures. A cryostat was used for serial sectioning of the central brain region (CCF x 4.5–9.5 mm) for each animal. Each series consisted of 24 coronal sections, each 20 µm thick with an even spacing of 200 µm between sections. Sections were distributed 4 per coverslip and fixed soon after. Three technical replicates were obtained for each series generated.

### Fixation

Before fixation, samples were allowed to sit for 5 minutes to ensure proper tissue adhesion to the coverslip. The sections were then fixed with 4% (v/v) paraformaldehyde (PFA, Electron Microscopy Sciences, 15714) in 1x PBS containing 3 mg/mL PVSA for 10 minutes, followed by three 5 minute washes in 1x PBSTw. Sections were stored in 70% ethanol at −20°C until further use.

### Permeabilization

Samples were stored for a minimum of 72 hours to a maximum of 4 months before preparation. Upon removal from storage conditions, samples were washed three times in 1x PBStw for 5 minutes at room temperature, and permeabilized in 1x PBS containing 1% (v/v) Triton X-100 (Sigma Aldrich, X100) and 3 mg/mL PVSA for 30 minutes at 47°C with constant agitation.

### MelphaX RNA modification

Following permeabilization, the samples were rinsed three times in 20 mM MOPS (pH 8) containing 0.1% (v/v) Tween-20 and 3 mg/mL PVSA at 47°C for 5 minutes each. After the final rinse, the samples were aspirated dry, and 100 µL of 0.5 mg/mL MelphaX diluted in MOPS buffer was pipetted directly onto tissue sections for RNA modification. To prevent evaporation, a small piece of parafilm was placed over the MelphaX solution, and the samples were incubated at 47°C for 1 hour. MelphaX was prepared according to EASI-FISH protocol ^5^.

### Hydrogel embedding

After RNA modification, samples were rinsed three times for 5 minutes in 1xTBS (2 mM TRIS 300 mM NaCl) with 0.1% (v/v) Tween-20 and 3 mg/mL PVSA (1xTBStw) at room temperature. A gel solution containing 3% 19:1 acrylamide:bis-acrylamide (Sigma Aldrich, A3449) in 1xTBS with 0.1% (v/v) Tween-20 and 3 mg/mL PVSA was prepared for sample hydrogel embedding. 3 mLs of gel solution were made per coverslip and split, with 2mLs containing 0.1% (v/v) N,N,N′,N′-Tetramethyl ethylenediamine (TEMED, Sigma Aldrich, T7024) and 1 mL containing 1% (w/v) Ammonium persulfate (APS, Sigma Aldrich, A3678). Gel solution containing TEMED was added to samples for 30 minutes under vacuum allowing the solution to penetrate into the tissue sections. After degassing, the 1 mL of gel solution containing APS was added to the sample and briefly mixed by pipetting. A pedestal containing a coverslip, treated with gel slick (Lonza, 50640) was inverted onto the sample to form a gel between the pedestal and the sample, the excess gel solution was aspirated off. Samples sat for 2 hrs allowing the gels to polymerize fully before being separated from the pedestal with a razor blade.

### Pre-Clearing

Samples were washed three times for 5 minutes with 1xTBStw at 47°C. A 2% (v/v) sodium dodecyl sulfate (SDS, Thermo Fisher Scientific, AM9820) solution in 1xTBS with 0.1% Tween-20, 3 mg/mL PVSA, and 1% (v/v) proteinase K (New England Biolabs, P8107S) was used to clear the sample. Digestion was carried out at 47°C with agitation for 18 hrs and then rinsed three times for 5 minutes with 1xTBStw. All steps in clearing, including the washes before and after, should be done at 47°C to avoid precipitation of SDS.

### Encoding

A 50% (v/v) formamide (Thermo Fisher Scientific, AM9344) solution in 1xTBS with 0.1% Tween-20 and 3 mg/mL PVSA was used to equilibrate the sample in a hybridization buffer for 10 minutes at 47°C. Each coverslip was aspirated dry and hybridized with 30uL of encoding solution containing 600 ug of RNA encoding probes (150 ug/section) in 50% (v/v) formamide with 10% (w/v) dextran sulfate (Sigma Aldrich, D6924) and 1xTBS and 0.1% Tween-20 and 3mg/mL PVSA. The encoding solution was pipetted directly onto the sample before being covered with a parafilm square. Hybridization of encoding probes was carried out for 18 hrs before being washed 4 times for 15 minutes at 47°C with agitation in 50% formamide in 1xTBS with 0.1% Tween-20 and 3 mg/mL PVSA. For MERFISH, encoding was done with 30% (v/v) formamide at 37°C since the probes are DNA.

### Post-Clearing and Hydrogel embedding

Samples were washed three times with 1xTBStw for 5 minutes at 47°C and cleared a second time to further reduce encoding probes that may have bound non-specifically. Clearing was done for 3 hrs, as described above, and then washed with 1xTBStw at 47°C. A second hydrogel was formed on the sample, as described above, to reduce any lifting caused by tissue clearing of dense brain regions. Sample was washed three times in 1xTBStw for 5 minutes and stored in a 10% formamide solution in 1xTBS with 0.1% Tween-20 and 3 mg/mL PVSA at 4°C

### Automated Data Collection Hardware & Software

We developed a custom fluidics system to enable continuous high-throughput imaging across six 40-mm coverslips, addressing limitations of commercial fluidics systems in multiwell, high-flow environments. Our system allows for simultaneous imaging and hybridization by alternating between two groups of wells, substantially reducing liquid handling time and maximizing throughput.

A chamber was designed with six 35 mm wells, creating a watertight seal with the coverslips while preserving imaging space. The wells were configured in a tight two-by-three arrangement, fitting in the stage adapter footprint, with four M3 screw holes surrounding each well. This profile was used to mill 2 mm stainless steel plates for compressing the chamber and providing a flat imaging plane. The edges of the chamber design were then inset by 0.5 cm to account for expansion once assembled. To create the chambers, a negative mold was designed and 3D printed using PETG. Two part silicone was mixed and degassed for 15 minutes to minimize large bubbles in the chamber. The silicone was then poured into the mold and allowed to set overnight at room temperature.

The chamber is assembled by aligning a steel plate with a silicone chamber placed on top of it. Coverslips are then placed on top of the silicone chamber with samples facing the steel plate. A second steel plate is then placed on top of the samples and secured to the chamber using four M3 × 16 mm screws around each sample. The bottom of each sample is then cleaned using lens cleaner and lens paper, and a small sticker was added to the edge of each coverslip to serve as a reference point for autofocus. Once placed on the microscope, fluidics is integrated using a 3D printed lid, holding blunt-tipped needles in each chamber at an angle to prevent vacuum formation. This open-well design supports milliliters-per-second flow rates, substantially improving throughput compared to traditional closed chambers.

To accomplish milliliters-per-second flow rates a fluidics system was assembled consisting of a syringe pump, multiposition valves and python software control allowing automated control of up to 30 readout hybridization rounds and corresponding stripping steps on 6 large sample wells. A 5mL glass syringe was used to minimize system maintenance and improve system reliability at high flow rates. Vici 10-24 port multiposition valves were daisy chained to enable easy expansion to large numbers of readout solutions. Fluidics tubing ranged from 1/16 inch in paths that were shared across multiple solutions and 1/8 inch diameter for tubing that goes directly to solutions allowing for faster flow rates. Additional features including vacuum aspiration for faster liquid removal and syringe mixing within open wells were added to improve reproducibility.

Custom python software (https://github.com/wollmanlab/Fluidics) integrates the fluidics system with ATLAS protocols, featuring a modular, file-based control system compatible with microscope software control. The user-friendly GUI allows for real-time simulation of protocols and manual control, facilitating flexible protocol development and execution.

### Chamber preparation for data acquisition

Fluidics system was cleaned by flushing with ethanol and rinsed with 1xTBStw before each experiment and stored in Ethanol for longer durations. Deep cleaning was performed by flushing with 10% Bleach before standard cleaning if contamination was suspected. Up to 6 coverslips were assembled into each chamber. Samples were stained with 1xTBS with 0.1% (v/v) Tween-20 and 3 mg/mL PVSA with 2ug/mL 4’,6-diamidino-2-phenylindole (DAPI, Sigma Aldrich, D9542) for 5-10 minutes at room temperature manually and rinsed three times with 1xTBS with 0.1% (v/v) Tween-20 and 3 mg/mL PVSA before being attached to the fluidics system.

### Data Collection

Images were captured on a custom Epifluorescent microscope with a 10X/0.45 NA Objective. Excitation light was provided by Solis LEDS for imaging Cy5 disulfide conjugated readout probes and PCB mounted LEDs for imaging dapi for nuclear stain. Emission was collected on FLIR Blackfly USB Camera with a pixel size of 0.425-0.49 µm. Microscope was controlled via Micromanager and custom MATLAB interface ^6^.

### Focus and Position Selection

Initial focus was set manually and entire coverslips were imaged to visualize dapi. Positions containing sections were manually selected using a custom drawing script. 5 evenly spaced positions were manually focused for each section as well as a reference position containing a registration sticker per coverslip. A plane was fit for these positions per section to extrapolate focus and set relative to the registration sticker focus. Before each round of imaging the registration sticker was imaged and the focus plane was adjusted to ensure cells were in focus across the multiple days of staining, stripping and imaging.

### Automated Strip

Samples were stripped from fluorophores similar to MERFISH protocol ^4^ and imaged before the hybridization of each new readout probe, allowing them to be used as a background image for downstream analysis. Fluorophores attached with a disulfide to readout probes were stripped from the sample with 2.5 mL of 0.25 mM Tris(2-carboxyethyl)phosphine hydrochloride (TCEP, AKSci, X4741) in 1xTBS with 0.1% (v/v) Tween-20 and 3 mg/mL PVSA. Samples were incubated in TCEP solution at room temperature for 30 minutes, and mixed once halfway through before being rinsed three times with 1xTBStw before imaging.

### Automated Hybridization

Samples were briefly rinsed once in 30% (v/v) formamide in 1xTBS with 0.1% (v/v) Tween-20 and 3 mg/mL PVSA with 2ug/mL 4’,6-diamidino-2-phenylindole (DAPI, Sigma Aldrich, D9542). Readout probes, diluted to 10 nM in 30% (v/v) formamide in 1xTBS with 0.1% (v/v) Tween-20 and 3 mg/mL PVSA and 2ug/mL DAPI, were added to the sample. Hybridization of readout probes was carried out for 30 minutes at room temperature, with one mixing step half way through. Excess readout probes not bound were washed from the tissue three times with 30% (v/v) formamide in 1xTBStw with 2ug/mL DAPI, and twice with 1xTBStw, before imaging.

### Image Processing

Raw images were processed using custom python code available in the project’s Github repo (https://github.com/wollmanlab/ATLAS_Mouse_Brain/). In short, raw images were binned to a pixel size of 0.85-0.98 µm. Camera constant and excitation light contamination was experimentally calculated for each acquisition and subtracted. Uneven illumination and emission capture was experimentally calculated for each acquisition and corrected. Images were subjected to a 2 pixel median low pass filter to remove hot or dead pixels as well as a 25 pixel sigma rolling ball high pass filter to remove residual constant as well as background that is much larger than cells.

Background acquisitions were registered to readout acquisition using cross correlation on dapi images. Background acquisitions were then subtracted from readout acquisitions for only the signal channel. Processed images were then registered to a reference hybridization round and stitched together using cross correlation of dapi images.

Cells were segmented using Cellpose ^7^ cyto3 model on the stitched reference dapi image for nuclei as well as a max projection of all readout images for total cell. Missed cells in dense areas were recovered by calling peaks in dapi images and morphologically dilating to a 5 µm radius. For each cell the median of the segmented pixels for each measurement was calculated and saved for future analysis as well as the cells segmentation properties and spatial coordinates.

### Common Coordinate Framework Registration

Each section was registered to the common coordinate framework by first assigning an approximate CCF x through visual inspection with reference MERFISH data. Registration in z and y were performed by manually clicking registration points and fitting a radial basis function model to convert from experimental spatial coordinates to CCF coordinates.

### Low Quality Cell Filtering

Non cells were removed as cells below a sum signal threshold as well as beyond 50 µm from a connected component graph created from the spatial coordinates of all cells. Low Quality cells due to hydrogel integrity or registration errors were removed as any cell that had a dapi signal decrease of more than 50% in 2 or more rounds.

### Cell Scalar Correction

To correct for uneven staining and total RNA content the overall magnitude of each cell vector was normalized. First a robust magnitude was approximated by correcting bits where cells were outliers and then taking the sum of the corrected cell vector. Cells were normalized by scaling their approximate magnitudes to the same value. Residual scalar differences due to the position of each cell in the optical field of view were corrected by fitting a linear regression between image coordinates and each measurement.

### ATLAS vector normalization

Measured cell vectors were normalized independently for each latent dimension by subtracting the median signal across cells and scaling by the robust standard deviation estimated from the 1st to 99th percentile range; this linear normalization was used to reduce batch-to-batch differences in latent-space amplitude before clustering or decoding.

### Unsupervised Clustering

An igraph implementation of Leiden clustering was then performed with a high-resolution parameter. Clusters within correlation of 0.9 were then merged.

### Decoding & Harmonization

To decode ATLAS signatures into predefined cell types, we employed a Bayesian recursive harmonization and classification approach called SCALE (Single Cell Alignment Leveraging Existing data). This method maximizes the use of reference atlas data and incorporates spatial priors derived from the anatomical structure of the brain.

Reference ATLAS Vectors: Reference ATLAS vectors were computed by projecting scRNAseq data (main text ref 4) through the DPNMF projection matrix, reducing the data to an 18-dimensional space. This dimensionality reduction enabled us to align the scRNAseq data with spatial transcriptomic signatures measured by ATLAS.

Spatial Priors: Spatial priors for each cell were generated using a kernel density estimate of cell type distributions based on CCF-registered reference MERFISH data. The kernel density was approximated by using numpy histogramdd on the ccf coordinates for each type with a binsize of 100 µm. The three dimensional histogram was then smoothed using a gaussian filter with a sigma of 100 µm in the CCF y and z axes and 250 µm in the x-axis. To calculate the spatial prior of a single cell, the ccf coordinates of that cell were used to pull the density estimate of each cell type in that location, normalizing to a sum of 1. These spatial priors represent the probability of finding specific cell types in certain brain regions, leveraging known anatomical structures. Spatial probability maps for each subclass were constructed across the brain using these priors.

Balanced Reference Vectors: To ensure an even comparison across sections, reference vectors were sampled based on the average spatial priors of all cells in a given section, creating a section-balanced reference. Reference vectors and measured vectors were normalized using robust magnitude and bitwise corrections.

Initial Neuron vs Non-Neuron Classification: Unsupervised clusters using cellular transcriptional signatures were classified into neurons and non-neurons using a pynndescent based KNN classifier trained on section-balanced reference cells with Euclidean distance as the metric and 25 neighbors. After this neuron/non-neuron classification, the spatial priors were updated to include either only neurons or only non-neurons, depending on the classification of the cluster.

Recursive Subclass Classification: For subclass-level classification, we employed a recursive decision tree method. The decision tree structure was constructed by averaging the cell type vectors from reference scRNAseq data and fitting a dendrogram based on pairwise correlations. Cell type average vectors can be found in Supplementary Table 4. At each binary decision point in the tree, two operations were performed:

1. Posterior Probability Calculation: For each cell, the posterior probability of belonging to one of the two subtrees was calculated. This was done by combining the spatial prior with the likelihood of the cell type, inferred from a scikit-learn logistic regression classifier trained on section-balanced reference cells.
2. Harmonization: After each binary decision, the median and robust standard deviation (std) of the measured data were aligned with reference cells within the selected subtree. This linear correction ensured that the measured data was harmonized with the reference scRNAseq data.

Final Subclass Assignment and Imputation: Cells continued through this recursive decision process, moving down the decision tree and being harmonized at each step until they reached a final subclass label at the leaves of the tree. At this point, cell types were assigned, and the data was fully harmonized with the reference atlas. Imputation of transcriptomes was then performed using a nearest neighbors approach based on transcriptional signatures. Using the harmonized vectors for the measured cells the 15 nearest reference neighbors were calculated using pynndescent based KNN with Euclidean distance as the metric. Imputed expression was calculated as the average gene expression for these 15 reference cells. Subclass names, numeric identifiers and visualization colors were assigned in agreement with Allen whole mouse brain reference and can be found in Supplementary Table 1.

### Validation - pairing two datasets

The accuracy of imputation was calculated using pseudo-pairing procedure. Two matching sections from different datasets (e.g.. ATLAS imputed and MERSCOPE) based on CCF x values after registration done as described above. To account for overall sampling frequency the total number of cells were downsampled to the same number. Cells were then paired across datasets in a greedy pairing algorithm, as finding global optimal pairing was too computational time consuming. The greedy pairing first created a list of all candidate pairings of all pairs of cells 100 μm from each other. The correlation score for all candidate pairs was calculated based on the Pearson correlation of the log gene expression across the 336 shared genes among all brain atlases. The correlation scores were sorted from highest to lowest, and the greedy algorithm selected pairing according to this sorted list after each selection, excluding possible pairing that included a cell that was just paired. The output of this pairing algorithm was the correlation score between all optimally paired cells. The cumulative distribution function of these correlation scores was used to evaluate the quality of agreement between the two datasets. Higher agreement results in increased area under the curve and more cells above each threshold.

### Validation marker gene selection

The genes for MERFISH validation of subclass types were selected from a set of 500 genes in the referenced MERFISH data. Genes that contributed most to subclass classification were prioritized by selecting those with the highest F1 scores. To enhance the signal in MERFISH imaging, genes with more than 96 hybridization probes were chosen. The number of bits also aligned with the number of hybridization rounds used in the ATLAS data collection, while maintaining a Hamming-Distance-4 for error correction. 17 ‘blank’ barcodes were included to measure the false-positive rate in MERFISH measurement, resulting in a final selection of 170 genes.

### Validation marker gene MERFISH Imaging

Images were captured on a custom Epifluorescent microscope with a 63x Objective. The excitation light for imaging Cy5 was consistent with the ATLAS protocol. The PCB-mounted UV LED was used for imaging the fiducial markers and dapi. Emission was collected with a pixel size of 0.083 µm. For each position, 4 z-indexes were captured per dataset with 1 µm spacing. MERFISH imaging was done using FCS2 chambers as described ^4^.

### Validation marker gene - combining ATLAS and MERFISH analysis

After completing the 18 rounds of MERFISH hybridization and imaging, the FCS2 chamber was disassembled, and the coverslip with the tissue was placed in a petri dish. To remove the marker gene DNA encoding probes, the sample was washed 4 times for 15 minutes at 47°C with agitation in 50% formamide in 1xTBS with 0.1% Tween-20 and 3 mg/mL PVSA. The sample was hybridized with the ATLAS encoding probes following the previously described protocol. The fluidics system was cleaned as outlined earlier, and the microscope objective was switched to 10x. After the post-encoding clearing and washes, the sample was assembled back into the FCS2 chamber and imaged according to the ATLAS protocol.

### Validation marker gene MERFISH Image Analysis

Image analysis was performed using custom Python code (https://github.com/wollmanlab/PySpots). In short images were corrected by replacing hot pixels with the medians of their immediate neighbors before being subjected to a highpass and lowpass filter to remove background and high frequency noise. Readout rounds were registered to a reference hybridization round using fiducial markers. Spots were called with the trackpy locate function. Spots were paired across rounds of hybridization into candidate transcripts. Transcripts that matched designed barcodes within 1 error were assigned to cells using a segmentation mask generated with Cellpose on dapi images. Cells with fewer than 10 transcripts were removed. Individual fields of view were registered between MERFISH segmentation and ATLAS segmentation using a rigid transformation to minimize the residual distance between paired segmentation masks. Cells with paired masks that were larger than 2 μm were filtered to ensure accurate pairing between datasets.

### Construction of 3D volumetric cell type inference

To analyze the spatial distribution of cell types across mouse brain samples, we developed a data processing pipeline to convert individual cell coordinates and types into a standardized 5D tensor. For each of the 15 animals, the X, Y, Z coordinates and cell types were binned into a 100 µm grid in the tissue section plane (CCF z and CCF y) and 300 µm along the anterior-posterior (AP) axis, which accounts for the 200 µm sectioning performed. The AP axis was upsampled by a factor of 3 relative to the tissue section plane to increase resolution and create an isotropic matrix. This binning was implemented for each animal and cell type using NumPy’s histogramdd function.

A key challenge was managing incomplete data due to section tears and holes. Voxels with cell counts below 50% of the average cell count per voxel were replaced with missing values, encoded as NaN (not a number). To address unequal sampling densities across animals, a correction factor was applied based on the average number of cells per voxel across all samples, assuming that total cell counts per voxel remain constant across animals. A Gaussian filter with sigma of 150 µm was applied to the data to reduce noise and interpolate values in low-density regions. The resulting 4D tensors from each animal were then stacked to create a 5D tensor (sample, X, Y, Z, cell type).

### Comparative analysis - cell type spatial distributions

The statistical analysis of cell type distributions used the 5D tensors described above. Assuming brain symmetry, the tensors were reshaped so that the two hemispheres from each section were treated as separate samples, with one hemisphere flipped horizontally. For each comparison (male vs. female or B6 vs. BTBR), the relevant hemispheres were selected. For each cell type, the average counts at each voxel were calculated and rescaled to have the same mean, ensuring that the identified differences reflected spatial changes rather than global abundance.

A per-voxel 3D matrix of the differences between the two distributions was then computed. The residuals from this comparison were used to calculate the entropy of the spatial distributions. Residuals made the comparison spatially explicit, as voxel-wise abundance was compared across identical locations. Entropy was chosen to quantify the overall extent of the differences, with identical distributions yielding zero residuals and thus zero entropy, and highly distinct distributions producing high entropy.

To assess statistical significance, we applied a permutation procedure, where sample indices were permuted before averaging for each cell type. This allowed estimation of the probability distribution for the residual entropy. We used an adaptive sampling approach, performing an initial 1,000 permutations for each cell type and extending to 10,000 permutations when p-values were estimated to be smaller than 0.005. Significant cell types are reported in Supplementary Table 6.

### Comparative analysis - regional compositional analysis

Similar to the cell type analysis, the regional analysis used the inferred 5D tensors as input, assumed brain symmetry, and calculated the average 4D tensor (X, Y, Z, Type) for each condition (B6 male, B6 female, BTBR male). For each voxel, the correlation distance (1 - Pearson correlation coefficient) between the type distributions of the two conditions was calculated. This correlation distance was then compared to a null distribution generated through a permutation procedure, as described in the cell type analysis. We used a similar adaptive procedure for p-value estimation, starting with 1,000 permutations and extending to 10,000 when p-values were smaller than 0.005. To assess sensitivity to the spatial prior, we repeated SCALE decoding without spatial priors, recomputed the regional compositional p-value maps, and quantified agreement with the original analysis as the intersection-over-union of thresholded significant voxels across p-value thresholds.

### spDEG analysis

To identify genes whose expression patterns statistically and significantly deviate from a uniform distribution, we performed a spDEG a^n^alysis9. For a single coronal section, we first constructed a neighborhood graph where cells are nodes and edges are drawn between any two neighboring cells as defined by Delaunay triangulation. Then, for each gene, we calculated the average standard deviation between any cell and its 50 neighbors. This value was subsequently compared to the expected value of the average standard deviation from a randomly drawn sample of 50 cells from the population, disregarding their spatial location. A p-value was assigned by comparing the observed value to a permuted null distribution and corrected for multiple testing using the False Discovery Rate (FDR) for each gene and cell type combination. For this analysis, we only considered cell types with more than 1,000 cells in the section and genes with expression greater than zero in at least 100 cells. To compare ATLAS with direct spatial transcriptomic measurements, the analysis was restricted to genes shared with the MERSCOPE reference dataset and repeated on a matched MERSCOPE section using the same spatial-uniformity testing framework.

### Computational simulation of ATLAS performance

To estimate the expected accuracy of the ATLAS approach, a computational simulation was performed using 4 different single cell transcriptomics datasets. In total eight encoding designs were tested including the published 53 bit FISHnCHIPS ^10^ design and TreeDPNMF designs with varying numbers of bits [3,6,12,18,24,50,100 bits]. TreeDPNMF Encodings were designed using Smart-seq scRNAseq data as described in Encoding Matrix Design. To simulate ATLAS measured data, 1.4 million high confidence subclass labeled full transcriptome imputed data ^12^ generated from Allen Common Coordinate Framework registered MERFISH cells that were projected using the encodings as described in Decoding & Harmonization. For gene perturbation analysis, the gene expression of each cell and gene was changed by increasing or decreasing the value up to the gene perturbation percentage [0, 25, 50, 100 %] before projecting to simulate potential deviations between imputed gene expression and true gene expression within the cell. Simulated cells were decoded using SCALE as described in Decoding & Harmonization. Gene expression references used for decoding were generated by projecting Allen subclass labeled 10X scRNAseq data^11^ using the encodings as described in Decoding & Harmonization. Spatial prior used for decoding was generated using Allen subclass labeled cells collected on the MERSCOPE platform ^11^ registered to the Allen Common Coordinate Framework as described in Decoding & Harmonization. Accuracy was reported as the percentage of cells whose decoded cell type matched the imputed data cell type.

### Estimation of the effects of segmentation errors

To estimate sensitivity to segmentation errors, a representative measured section was subjected to perturbations of the Cellpose based segmentation masks. Perturbations consisted of either merging masks with their nearest neighbor cell, translational shift of the mask including the overlap of neighboring cells, erosion of the mask, or dilation of the mask including overlapping neighbor cells. Perturbed masks were then used to pull ATLAS vectors from measured images and decoded using SCALE as described in *Decoding & Harmonization.* Accuracy was reported as the percentage of perturbed cells whose decoded label matched the unperturbed Cellpose based mask. For merged cells, a cell was marked accurate if its decoded label matched either of the merged cells which is the equivalent of missing the other cell. Perturbations were performed in triplicate for a random 5,000 cells and for varying magnitudes for shift,dilation and erosion reported as a fraction of the average cell mask. To estimate the occurrence of errors in our Cellpose based masks, we manually annotated representative masks from Extended Figure 6a looking for cells whose masks were missed or split as well as cells whose masks were merged with another cell.

### Benchmarking gene imputation concordance

To estimate the reliability of gene imputation, six single-cell transcriptomic B6 mouse brain datasets across varying technologies were compared, including smFISH-based datasets [MERSCOPE, Zhuang MERFISH, Wollman MERFISH], imputation-based datasets [ATLAS imputation, Zhuang imputation], and sequencing-based data [10X]. To account for differences in capture efficiency across technologies, each cell’s gene-expression vector was normalized to a total expression of 1,000 before dataset-level averages were computed. Correlations were reported as the log log pearson correlation. Given that ATLAS Imputation was derived from 10X data, we compared the correlation of all smFISH based approaches to 10X to provide an expected agreement level between smFISH and sequencing. To account for brain sampling discrepancy, spatial datasets were filtered down to sections within the same CCF x window (7.8 mm) whereas sequencing datasets were sampled proportionally to the subclass abundance observed in the MERSCOPE sections. Correlation was performed on the 166 genes that all datasets shared. For a regional comparison, spatial datasets [MERSCOPE, Zhuang MERFISH, Zhuang Imputed and ATLAS Imputed] were compared as they had 544 shared genes. Regions were defined by the parcellation_structure of the MERSCOPE cells and only cells registered to that region were compared. Pairwise results were plotted as the percentage of regions for each pair above a threshold vs the pearson threshold. Three representative regions were shown for all pairs including ATLAS imputed as well as MERFISH vs MERSCOPE.

### Benchmarking spatial prior impact on performance

A key aspect of SCALE is the use of spatial priors to support the accuracy of harmonization between measured cells and reference cells. To estimate how sensitive the performance of SCALE is to the agreement between the spatial prior and the biology, we performed perturbations to the spatial prior and then performed simulation as described in Computational simulation of ATLAS performance. Perturbation was performed at various levels by performing dilation or erosion in three dimensions for each cell type. Accuracy was reported as the percent of cells whose decoded type matched the imputed data cell type.

### Quantifying the magnitude of anatomical agreement to B6

Registration to the Allen common coordinate framework consists of a rigid scaling, rotation, and translation as well as a non-rigid alignment. Animals with morphology consistent with the common coordinate framework are expected to have minimal difference between rigid and non-rigid alignment. To demonstrate this, for each cell the difference between rigid and non-rigid was calculated. A kernel density estimate for the average difference in millimeters was calculated for each 500 µm window between CCF x 4.5 and 9.5 mm with a voxel size of 50 µm. Areas with larger differences in BTBR mice than B6 mice indicate areas with lower agreement to the common coordinate framework and thus more anatomical differences.

### RNA-FISH validation of regional differences

To validate selected ATLAS-predicted regional composition changes, iterative RNA-FISH was performed in triplicate for male B6 and male BTBR brains across three sections containing identified spatial regions 4, 6, 8, and 10. The marker panel was selected to test regional abundance changes that could be evaluated with a compact RNA-FISH experiment, rather than to reproduce the full subclass-level ATLAS annotation. Slc17a6, Slc17a7, and Slc32a1 were included to distinguish major excitatory and inhibitory neuronal compartments, while Gja1, Sox10, and Adgrf5 were included to identify non-neuronal populations contributing to the predicted regional differences. Because these markers are not sufficient to define all neuronal subclasses across the whole brain, labels were assigned only within the preselected anatomical regions tested in Extended Data Fig. 15. Sections were prepared according to the described MERFISH protocol, with 40 encoding probes per gene and four readout arms per probe. Images were collected in 40X. Segmentation was performed on dapi with Cellpose plus a 5 µm dilation. Average intensity vectors for each cell were calculated for each round of RNAFISH. Background was removed at the image level by gaussian background subtraction and at the vector level by subtracting the 25th percentile across all cells to remove any residual background as no gene measured should be present in more than 75% of cells. To account for total RNA content and cell staining efficiency the vector for each cell was normalized to a sum of 1. To assign cell type labels, cells were filtered by region and then a global threshold across all sections was performed for each region based on marker genes according to Extended Figure 15c. For each section for each region the percentage of cells identified as a type was reported to account for differences in cell density and region capture size. Statistical significance was calculated using a student’s t test for each cell type and each region.

---

**Extended Figure 1:**
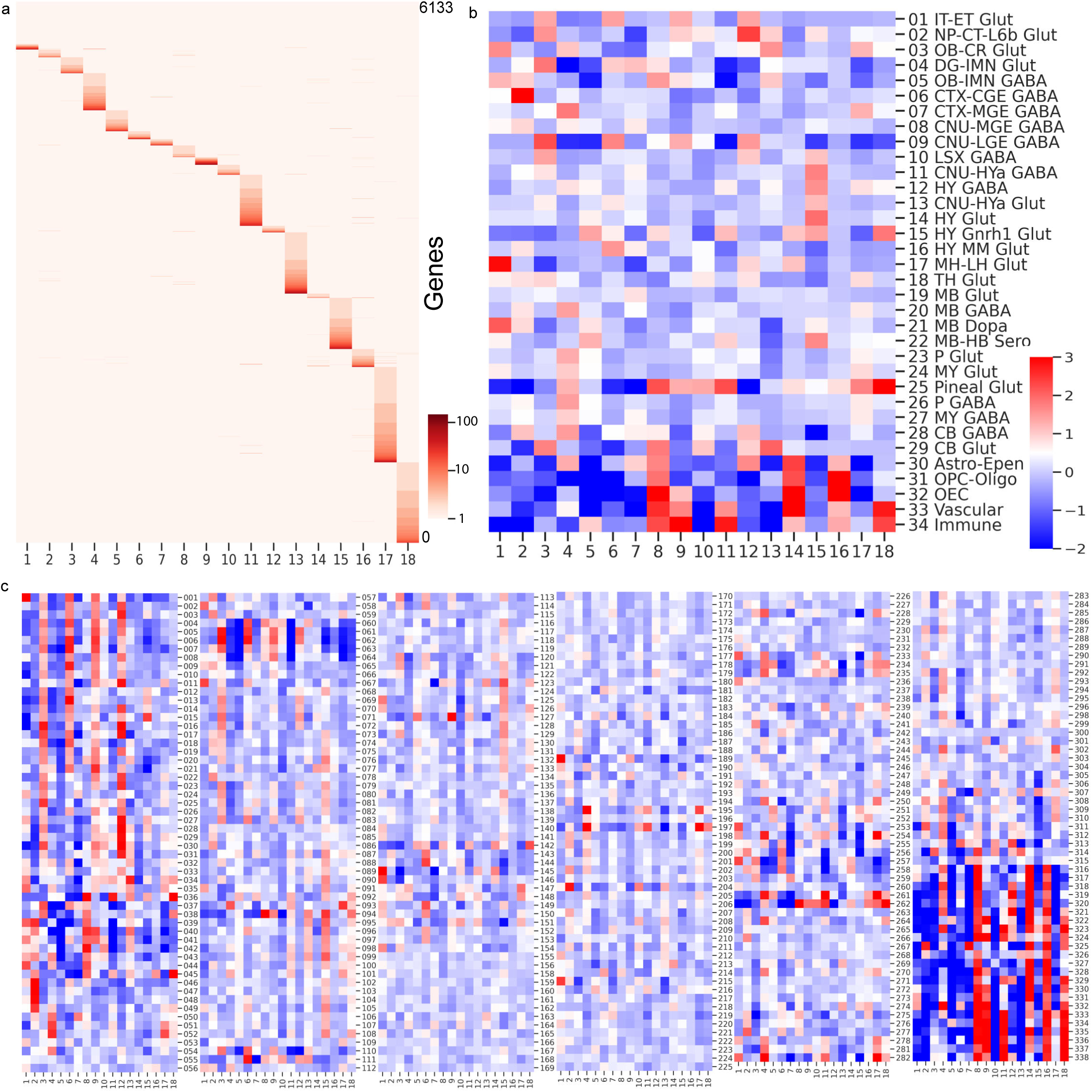
ATLAS DPNMF encoding design. **(a)** Heatmap of the DPNMF encoding matrix showing the log weight of each gene on each measurement. Weights directly translate into the number of encoding probes per gene per measurement. **(b)** Heatmap of the expected cell vector for each class-level cell type, calculated by projecting scRNA-seq data using the DPNMF encoding matrix and averaging across cell types. Values are z-score normalized. **(c)** Expected cell type vectors for all 334 subclass-level cell types. Values are z-score normalized.

**Extended Figure 2.**
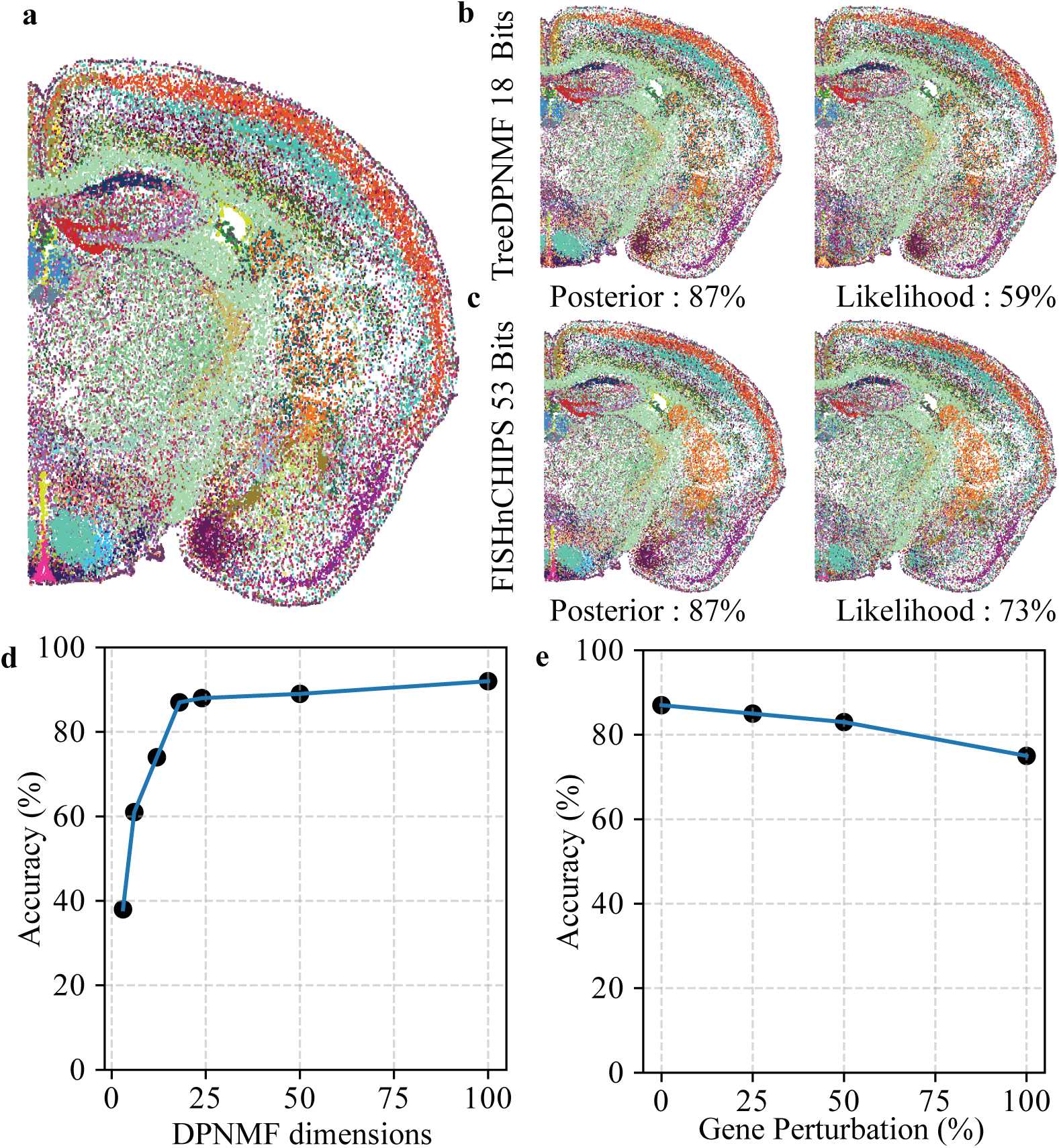
Simulation of ATLAS Performance on Imputed MERFISH Data. **(a)** A representative section from the MERFISH imputed dataset, which includes 1.4 million “High Quality Transfer” cells, is visualized with its assigned subclass labels. This dataset integrates anatomical coordinates (CCF), Allen subclass taxonomy labels, and a full transcriptome imputed from the original MERFISH data. **(b)** A simulation of ATLAS performance is shown using an 18-dimension TreeDPNMF projection. The decoding accuracy is 59% when based only on likelihood (without a spatial prior) and improves to 87% when a spatial prior is incorporated (Posterior). The reported accuracy represents the percentage of cells for which the decoded subclass label correctly matches the original Zhuang label. **(c)** A simulation of ATLAS performance is depicted using a 53-bit FISHnCHIPS projection matrix. Decoding without a spatial prior (Likelihood) yields a 73% accuracy, which increases to 87% with the inclusion of a spatial prior (Posterior). **(d)** Simulations of ATLAS decoding accuracy as a function of the number of TreeDPNMF dimensions used. Performance improves with increasing numbers of measurements, with diminishing returns after 18 measurements.

**Extended Figure 3:**
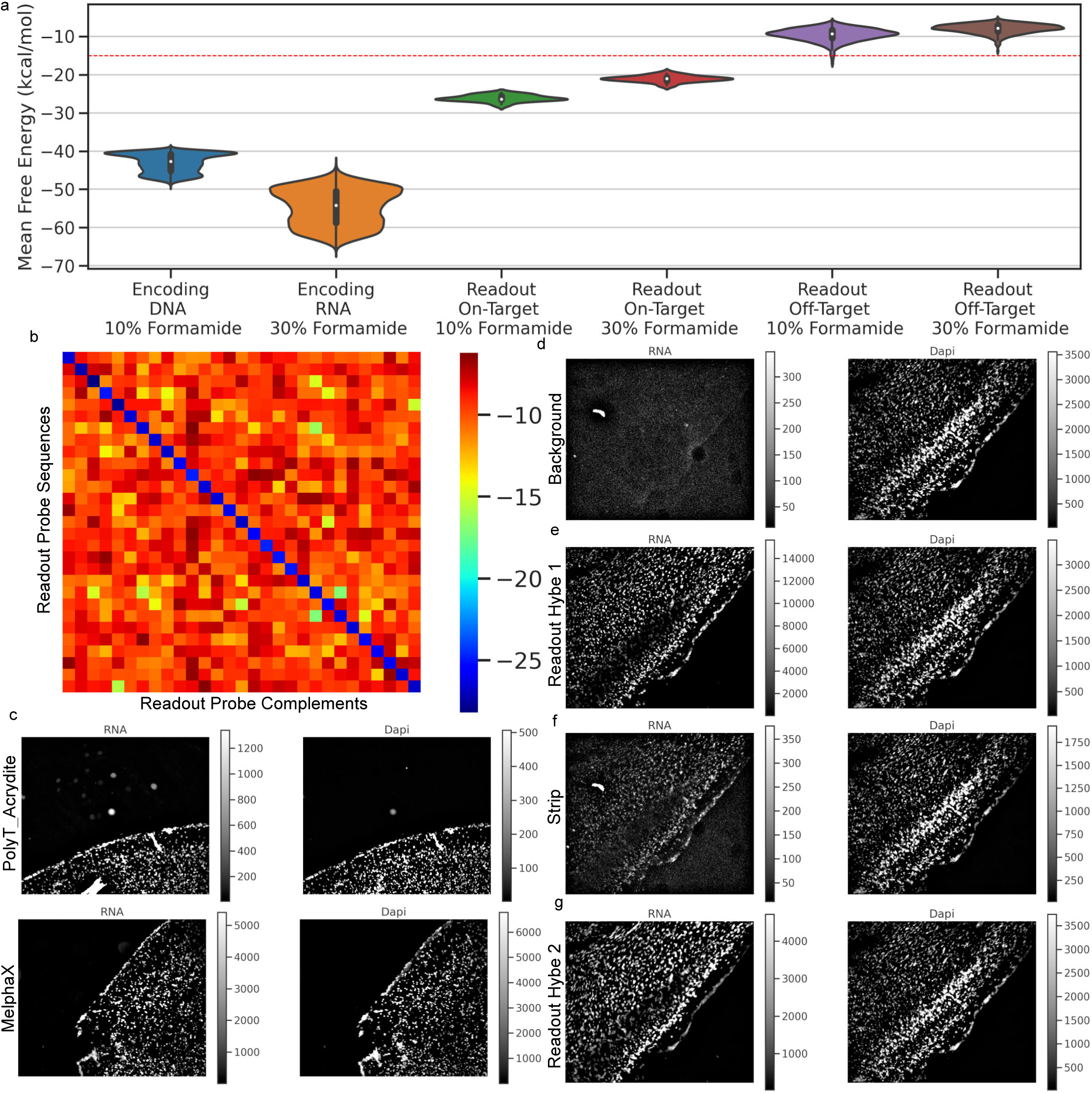
Experimental method development. **(a)** Violin plots of mean free energy calculations from NUPACK for hybridization event structures. Inset boxplots show the median (white point; 50th percentile), the interquartile range (thick bar; 25th–75th percentiles), and whiskers extending to the most extreme observations within 1.5 times the interquartile range. More extreme observations are not displayed separately. n=18 for readout ande 28,009 for encoding probes. **(b)** Pairwise heatmap of on- and off-target structure MFE stability scores. **(c)** Comparison of RNA and DNA signal between hydrogel anchoring strategies: PolyT-Acrydite, a hybridization-based approach, and MelphaX, a covalent approach. **(d)** Low background present in the RNA channel before readout hybridization. **(e)** Example signal observed with ATLAS signatures. **(f)** Residual signal remaining after TCEP reduction of disulfide-linked readout probes. **(g)** Subsequent readout hybridization, which contains signal from readout hybridization 2 as well as residual signal from readout hybridization 1.

**Extended Figure 4:**
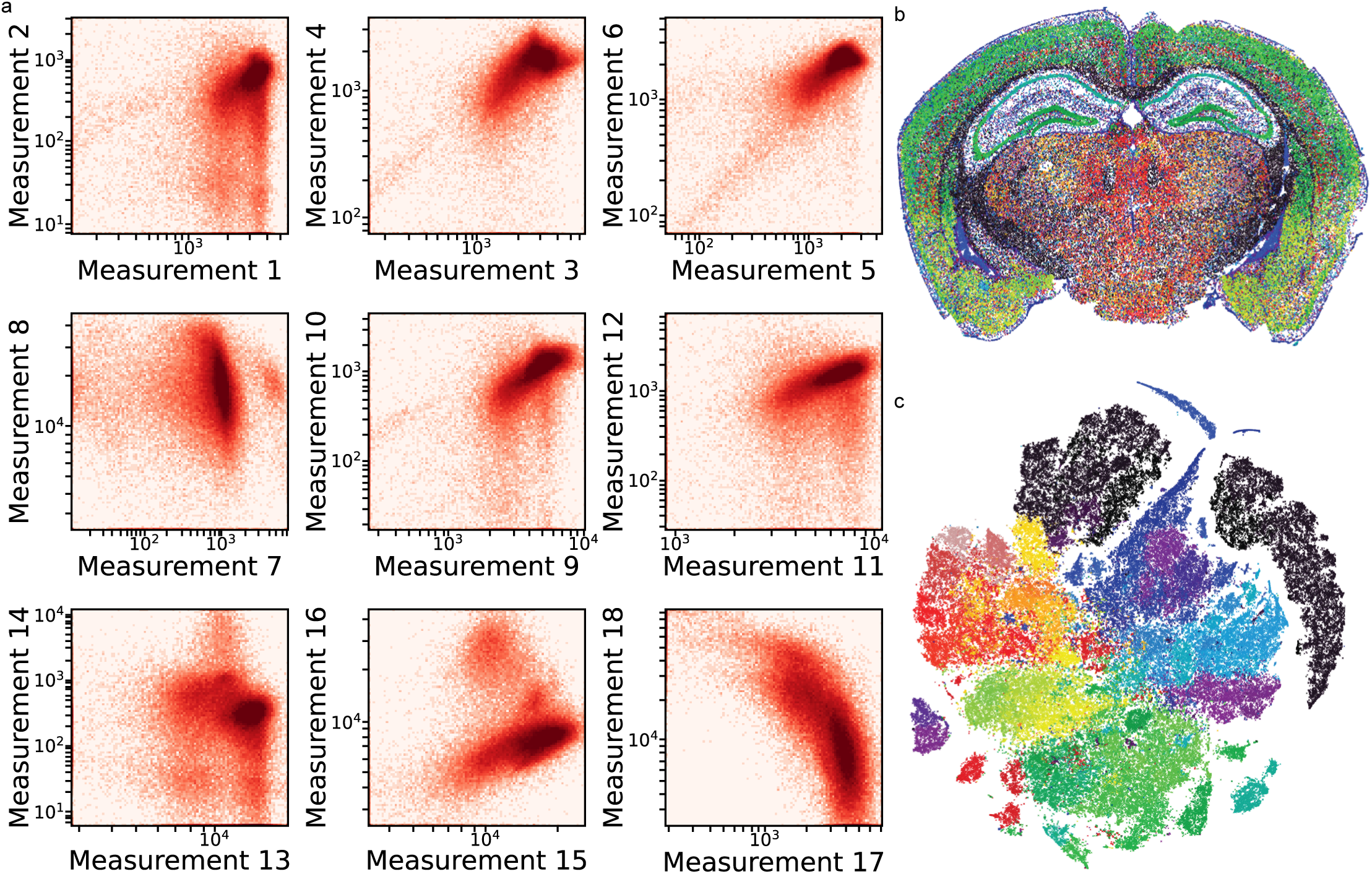
ATLAS transcriptional signature measurements. **(a)** Visualization of all 18 measurements for a single section. Multiple density modes are visible within each pairwise log10-scaled signal plot. **(b)** Spatial locations of unsupervised clusters from measured cells in a single section. **(c)** t-SNE representation of measured cells, colored by unsupervised Leiden cluster.

**Extended Figure 5:**
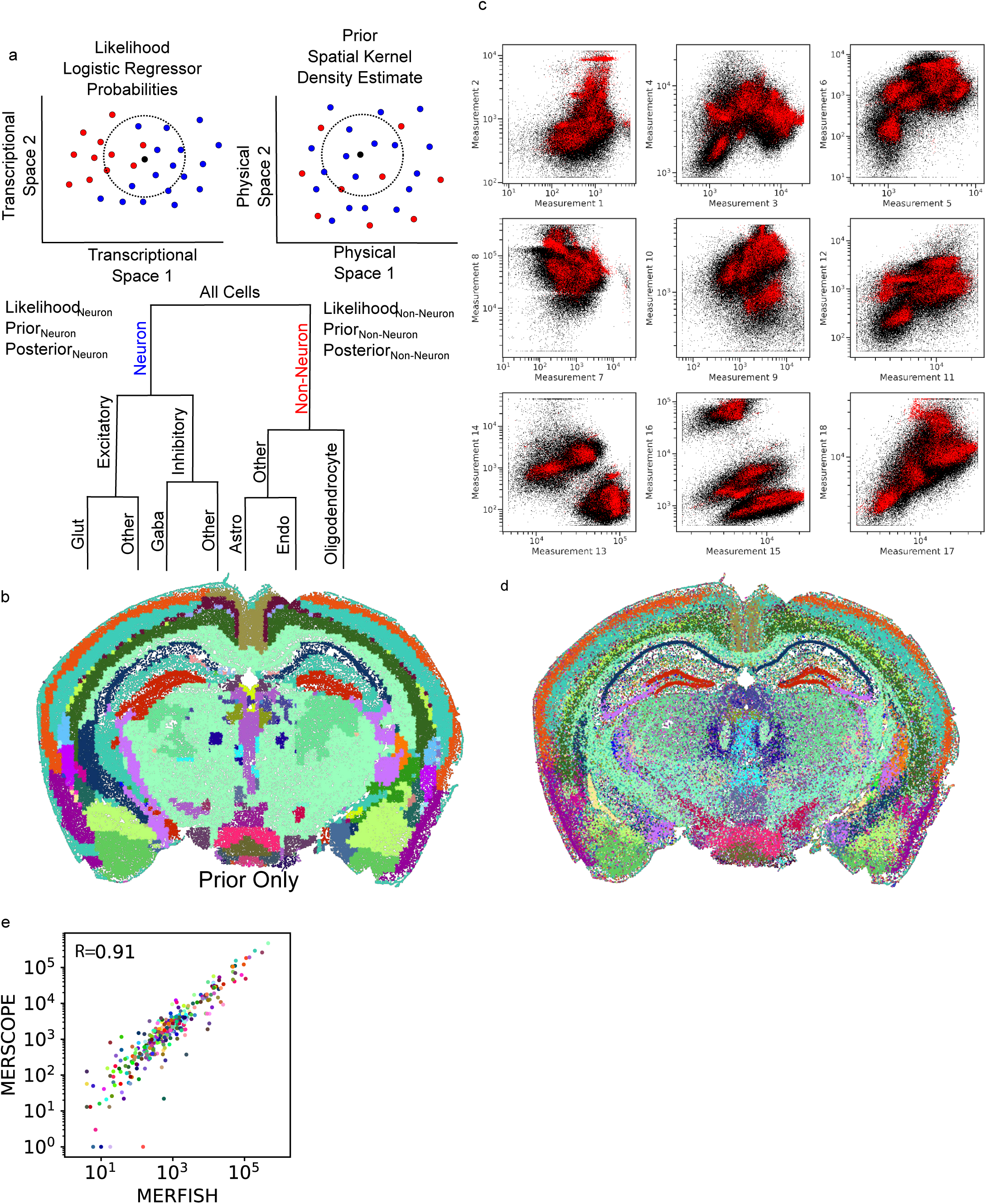
Decoding and Harmonizing. **(a)** Cartoon diagram showing the decoding strategy. Individual cells walk down a decision tree of cell types. Decisions are made by combining the likelihood that a cell belongs to a cell type, based on features calculated using a logistic regressor trained on reference scRNA-seq cells, with the spatial prior expectation of what cell types are present at that cell’s physical location, calculated using a kernel density estimate with CCF z- and y-axis sigmas of 100 µm and CCF x sigma of 250 µm on reference MERFISH data. **(b)** Visual representation of the spatial prior, demonstrated by passing measured cells through the decoder while ignoring likelihood. **(c)** Results of data harmonization. Cells are shown for each measurement, colored by source: black for reference scRNA-seq and red for harmonized measured ATLAS cells. **(d)** Results of data decoding at the subclass level. **(e)** Comparison of cell type abundance between MERFISH and MERSCOPE atlases. Each dot represents a cell type, colored using the BICCN convention (Supplementary Table 1).

**Extended Figure 6:**
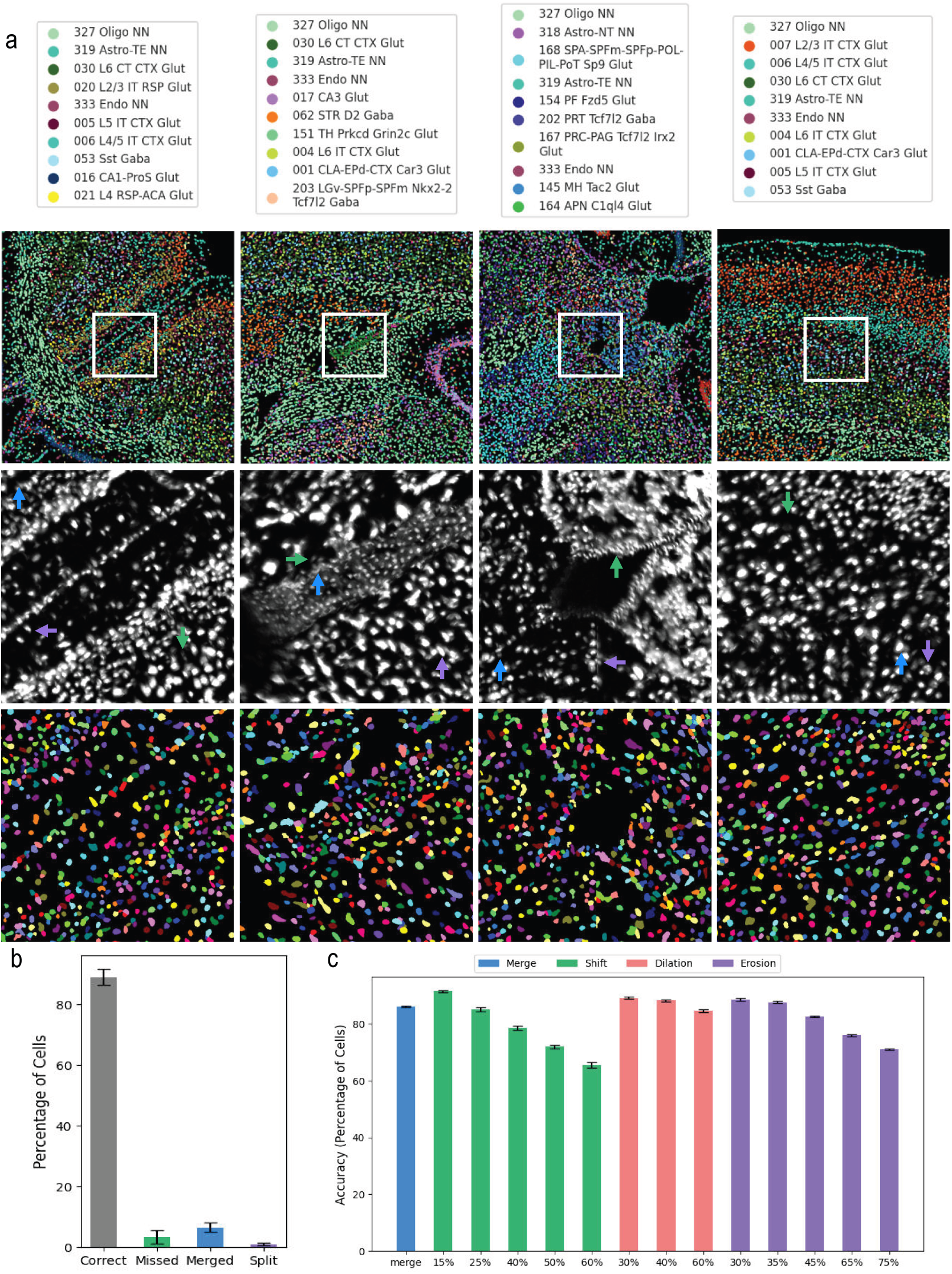
Cell Segmentation, Decoding Quality, and Simulated Impact of Segmentation Errors. **(a)** Representative images from four distinct brain regions illustrating overall SCALE decoding and Cellpose-based cell segmentation quality across diverse mouse brain regions. The top row shows brain regions color-coded by cell types. The middle row shows DAPI staining with arrows pointing to manually identified segmentation errors. The colors of the arrows correspond to the type of error as shown in panel b. Bottom row shows Cellpose segmentation. **(b)** Quantitative estimation of segmentation error rates. Manually scored segmentation in the four regions shown in panel a. **(c)** Simulation results demonstrating the impact of segmentation errors on cell type decoding accuracy. We simulated four types of errors: (i) merge, merging two nearby cells; (ii) shift, shifting masks by a percentage of cell area in a random direction; (iii) dilation, scaling masks by a percentage of cell area; and (iv) erosion, scaling masks by a percentage of cell area. In all cases, we updated the measured ATLAS transcriptional signatures based on the new segmentation masks and reapplied SCALE decoding. The reported accuracy is the percentageof cells that maintained the same label as before the synthetically introduced errors. The results show that while cell segmentation errors impact SCALE decoding, the decoding is still robust to errors of up to ∼30% incorrect masks. In panels b and c bars show average and error bars represent standard deviation across evaluated fields. n=30 FOV in panels b and c

**Extended Figure 7:**
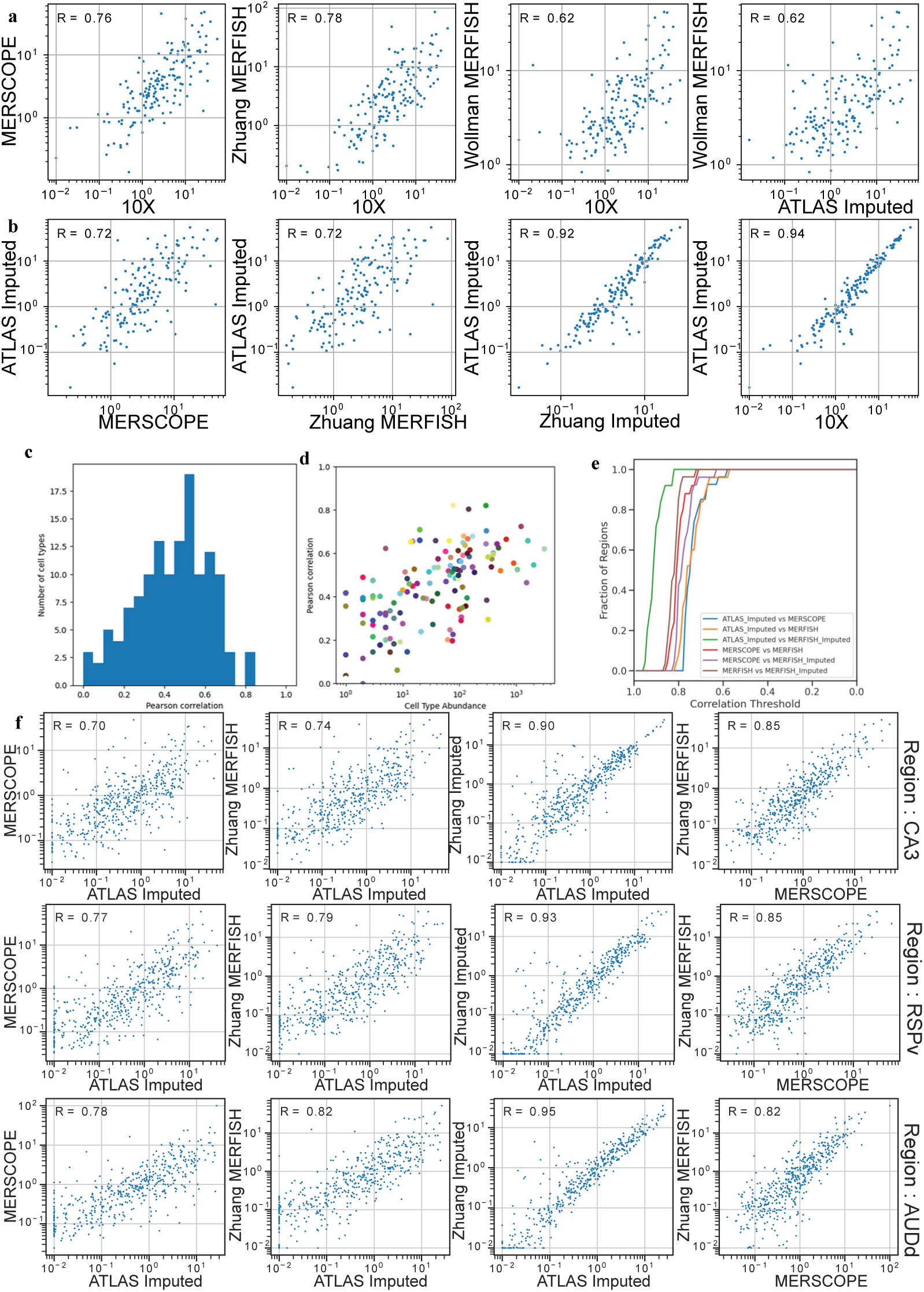
Benchmarking ATLAS Gene Imputation and Concordance with MERFISH Data. **(a)** Scatter plots comparing average gene expression on a log scale establish baseline correlations between different technologies and a 10X scRNA-seq reference to assess cross-technology concordance and ATLAS imputation accuracy. Comparisons show the correlation between 10X and a commercial MERSCOPE dataset, 10X and a published Zhuang MERFISH dataset, 10X and the Wollman lab’s MERFISH data, and a direct comparison of the Wollman lab’s MERFISH measurements against ATLAS-imputed expression from the identical cells. The correlation achieved by ATLAS imputation is comparable to leading spatial transcriptomics methods when compared to both scRNA-seq and direct MERFISH measurements. **(b)** Comparisons of ATLAS-imputed data show its correlation with MERSCOPE, Zhuang MERFISH, Zhuang-imputed data, and 10X data. **(c)** Analysis of gene expression correlation at the single-cell level, shown as a histogram of the distribution of Pearson correlation coefficients calculated between the 166 measured MERFISH marker genes and their corresponding ATLAS-imputed expression values. **(d)** A scatter plot relates cell type abundance to the median single-cell correlation coefficient for each identified cell type, exploring whether imputation accuracy is higher for more abundant cell types. **(e)** A cumulative distribution function (CDF) shows a paired correlation analysis using CCF regions rather than cell pairing. **(f)** Examples of gene expression correlations between technologies are shown for specific brain regions, including CA3, RSPv, and AUDd; each dot represents the average expression of an individual gene shared across the datasets.

**Extended Figure 8:**
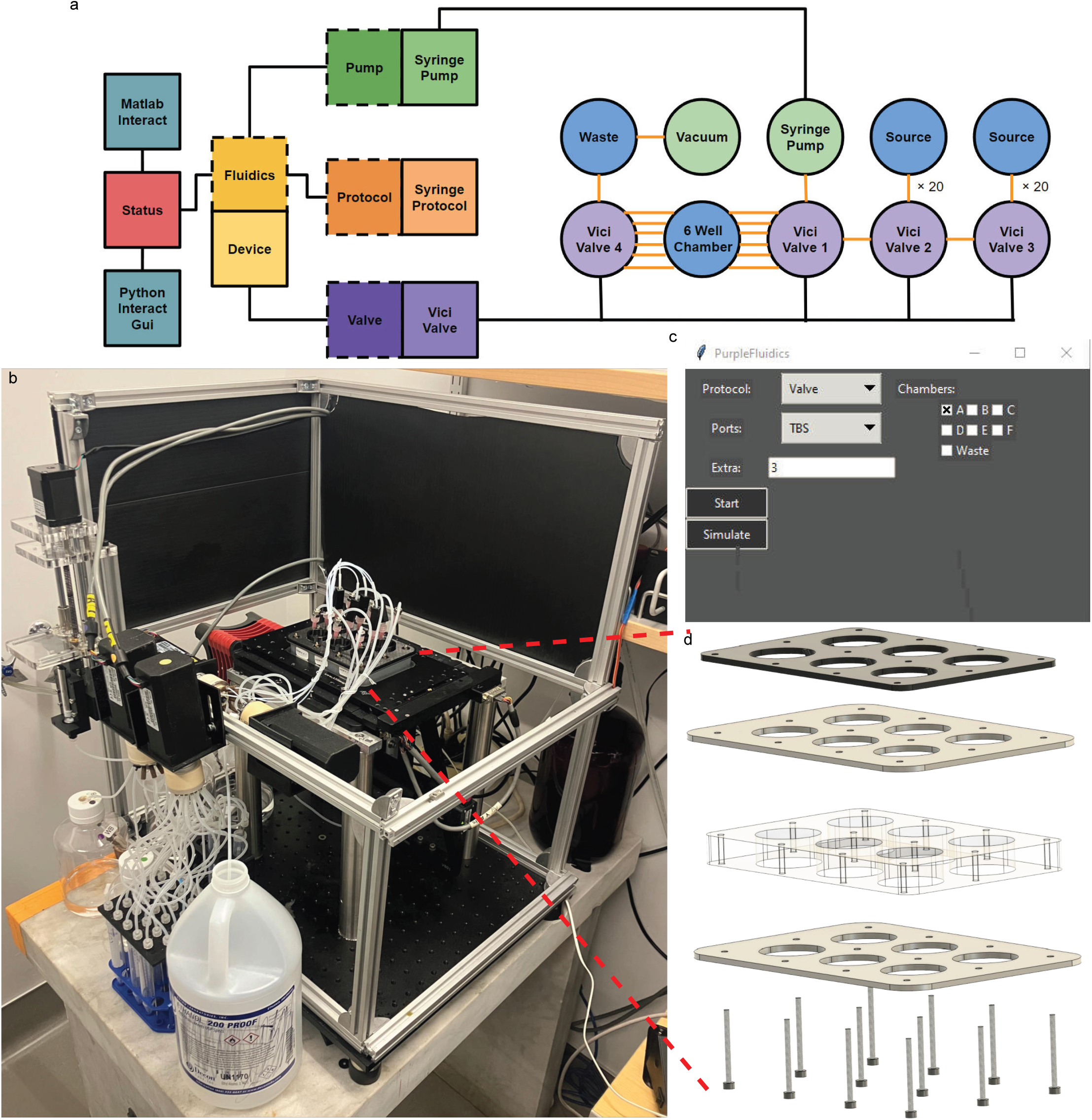
Automated Open-Chamber Fluidics System. **(a)** Schematic design of the Fluidics software architecture (squares) and hardware components (circles). Liquid is extracted from source tubes using an automated syringe pump and valves and deposited on sample coverslips. Waste is removed in reverse. Object-oriented software implementation allows modular integration of multiple device drivers, as well as multiple units of each device. The current implementation allows up to 30 rounds of readout hybridization and 30 rounds of stripping across each of six experimental wells. **(b)** Picture of the custom epifluorescent microscope with the open-source fluidics system attached. **(c)** Screenshot of the graphical user interface, which allows manual control when software control is not needed. **(d)** The open-well chamber design consists of six coverslips and a thick, soft silicone chamber sandwiched between two rigid steel plates. Two needles are placed in each well: one for aspirating liquid and one for adding liquid. The open chamber allows liquid exchange rates higher than closed systems, and the multi-well format allows simultaneous imaging and hybridization.

**Extended Figure 9:**
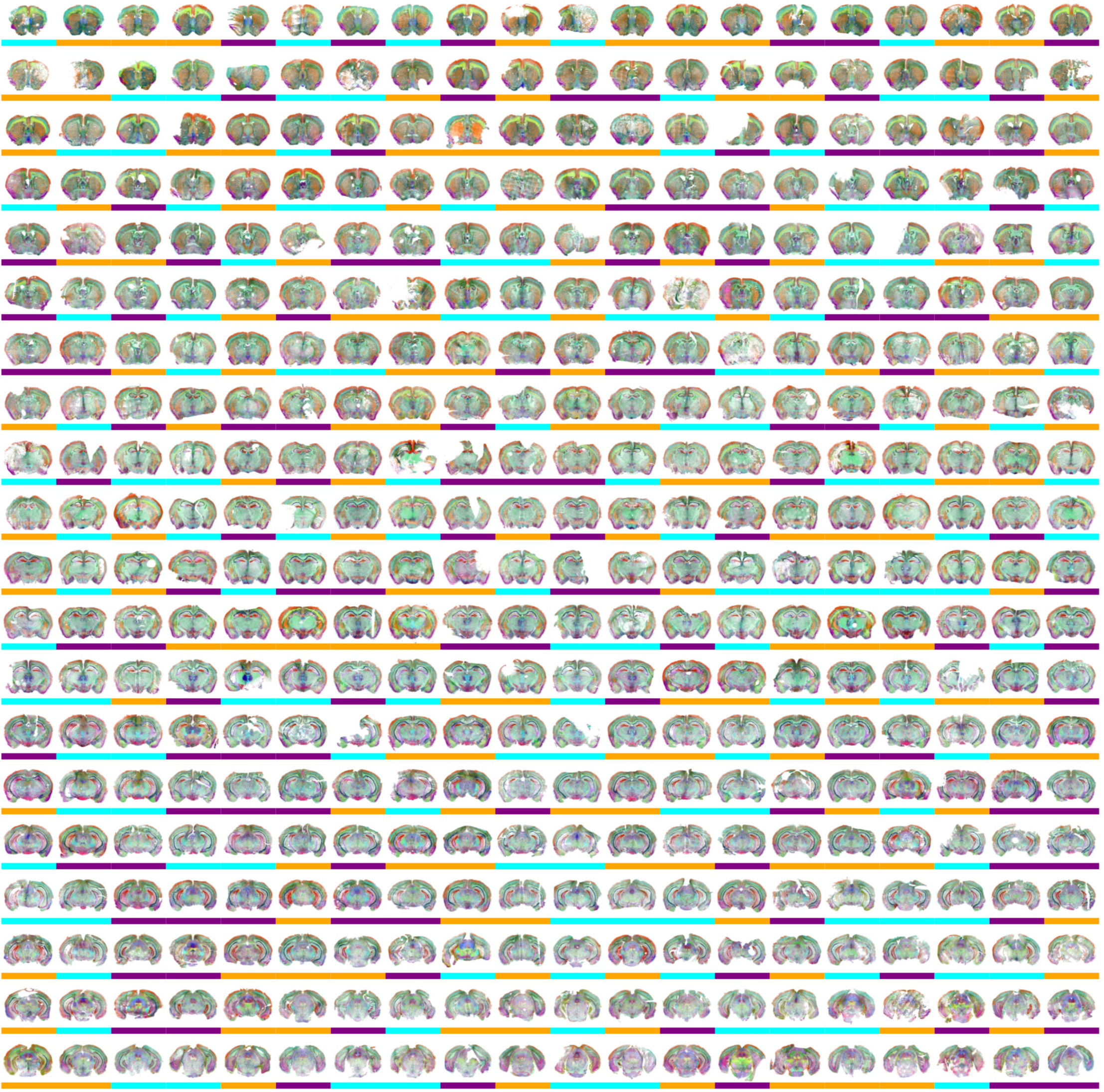
Dataset. Visualization of 400 sections collected across 15 animals spanning three genetic backgrounds: B6 male in orange, B6 female in purple, and BTBR male in cyan. Sections are ordered from anterior to posterior.

**Extended Figure 15:**
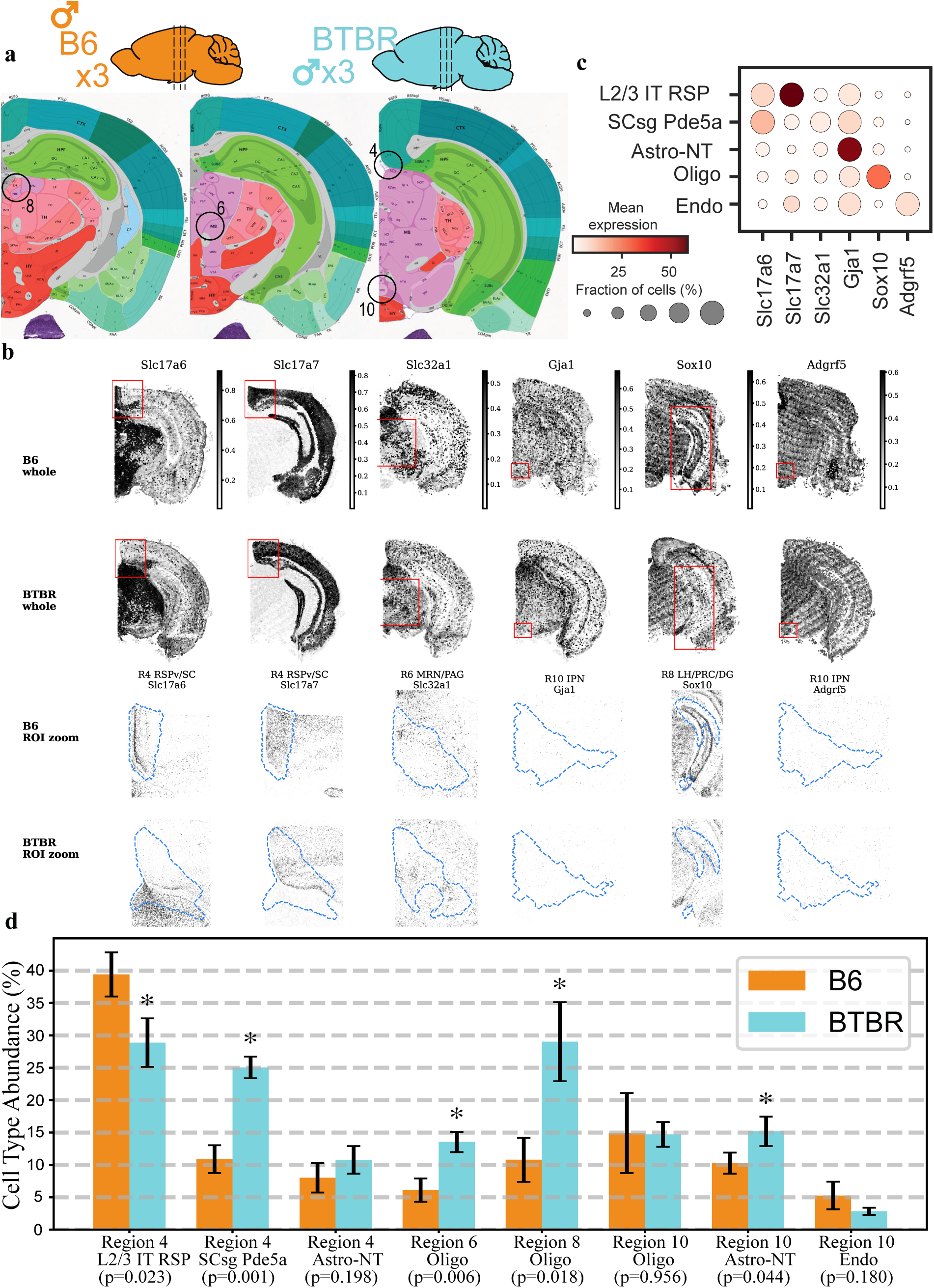
Experimental validation of ATLAS findings. **(a)** A diagram of the experimental design used to validate findings from the Autism Spectrum Disorder (ASD) model. Brains from wild-type B6 mice and the BTBR ASD model mice were sectioned, with overlays indicating the specific regions (4, 6, 8, and 10) selected for validation. **(b)** Representative RNA-FISH images for the six marker genes used in the validation. For each gene, whole-section images are shown for B6 and BTBR, with the red box indicating the region of interest used for the corresponding validation comparison. Matched ROI zooms are shown below for B6 and BTBR. CCF-defined region of interest are shown in light blue. **(c)** A dot plot showing the expression levels of the six marker genes across key cell types. The size of each dot indicates the fraction of cells expressing the gene, while the color indicates the mean expression level. **(d)** The bar plot compares the abundance of specific cell types in B6 and BTBR mice within the selected regions. Asterisks (*) indicate statistically significant differences (p-value < 0.05, two sided student t-test, n=20 hemispheres). Bars show average and error bars represent standard deviation across animals.

**Supplementary Figure 1.**
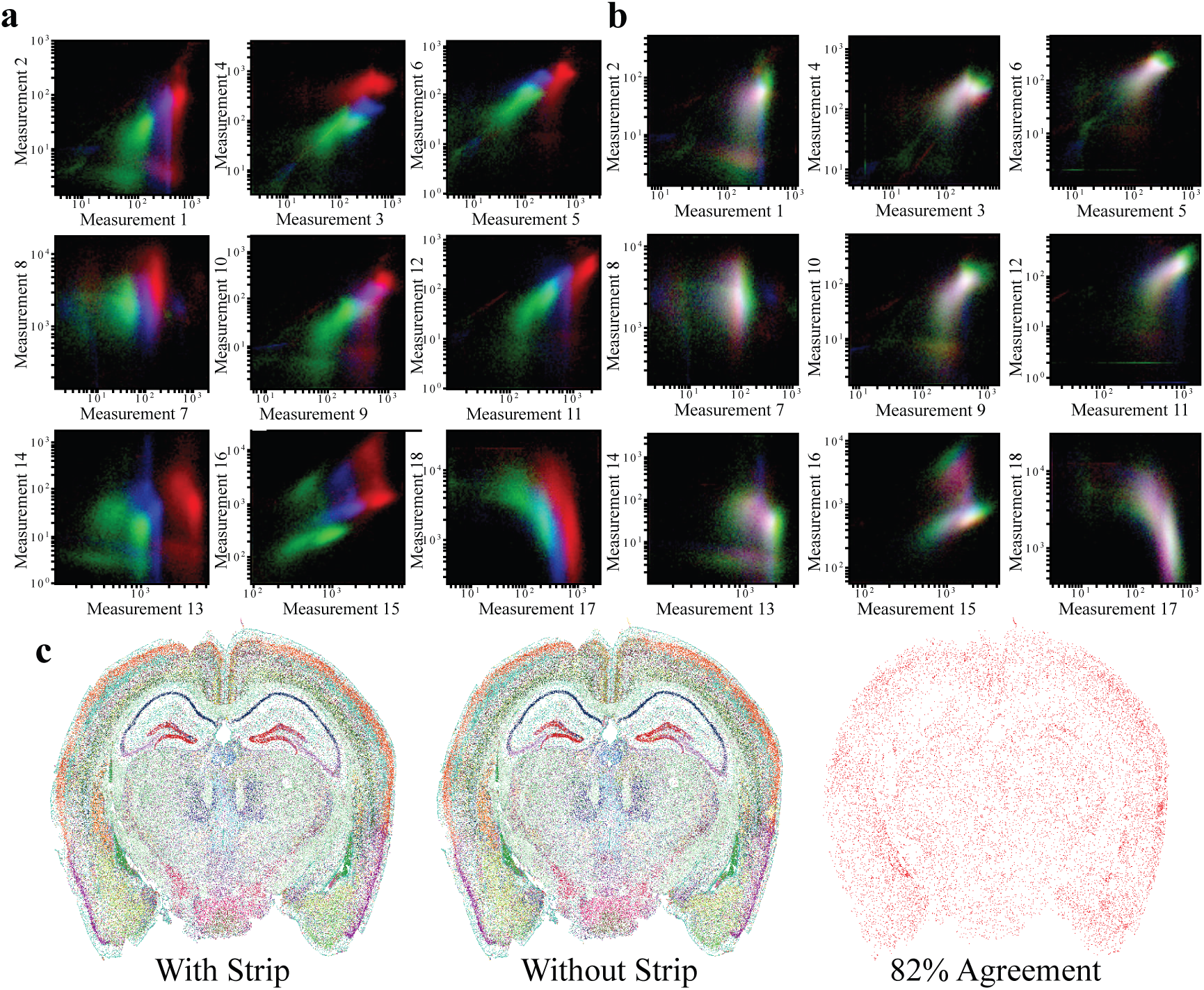
ATLAS latent-space measurement reproducibility and importance of background subtraction. **(a-b)** Visualization of the reproducibility of ATLAS latent-space measurements across three biological replicates, shown before and after linear rescaling. The data from the three replicates are represented by red, green, and blue color channels in correlation plots of the 18-dimensional latent-space vectors. **(a)**, The raw latent-space measurements show variability between the replicates, indicated by the separation of colors. **(b)**, After applying a linear rescaling to each replicate, the measurements demonstrate a high degree of overlap, indicated by the white color where the red, green, and blue channels mix. This shows that the underlying biological signatures are highly reproducible once systematic scalar differences are removed. **(c)**, Demonstration of the importance of background subtraction. The image on the left is a representative result that includes a background subtraction step (“With Strip”), while the middle image is a result generated without this step (“Without Strip”). The panel on the right indicates that there is an 82% cell-wise agreement between the results of the two methods.

**Supplementary Figure 2.**
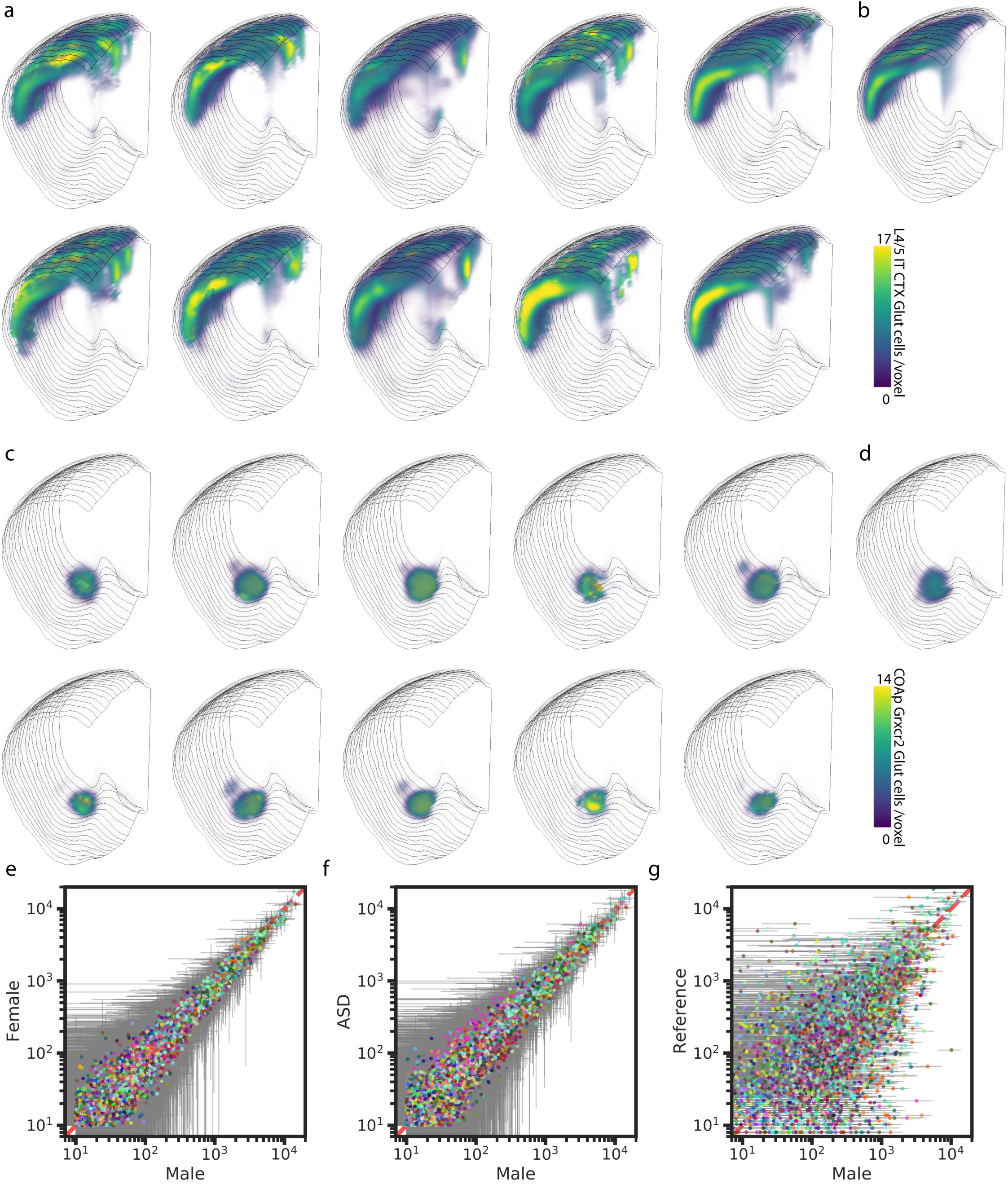
Reproducibility of spatial cell type abundance measurements. Panels **(a)** and **(b)** show the spatial distribution of L4/5 IT CTX Glut neurons, while panels **(c)** and **(d)** show COAp Grxcr2 Glut neurons. In (a) and (c), each subplot represents data from one of the ten hemispheres profiled using ATLAS. Panels (b) and (d) show corresponding distributions from the BICCN MERSCOPE reference dataset. Panels **(e-g)** compare cell type abundances across CCF regions between pairs of conditions using scatter plots: (e) female versus male; (f) BTBR versus male; and (g) MERSCOPE reference versus male. Each point represents a CCF region-cell type pair and is colored by the BICCN color code corresponding to the cell type identity. Error bars indicate the standard deviation across biological replicates **(n=l** O hemispheres per condition). No error bars are shown for the reference in (g), because the MERSCOPE dataset is derived from a singl animal. Erorbars show standard deviation across biological replicas.

**Supplementary Figure 3:**
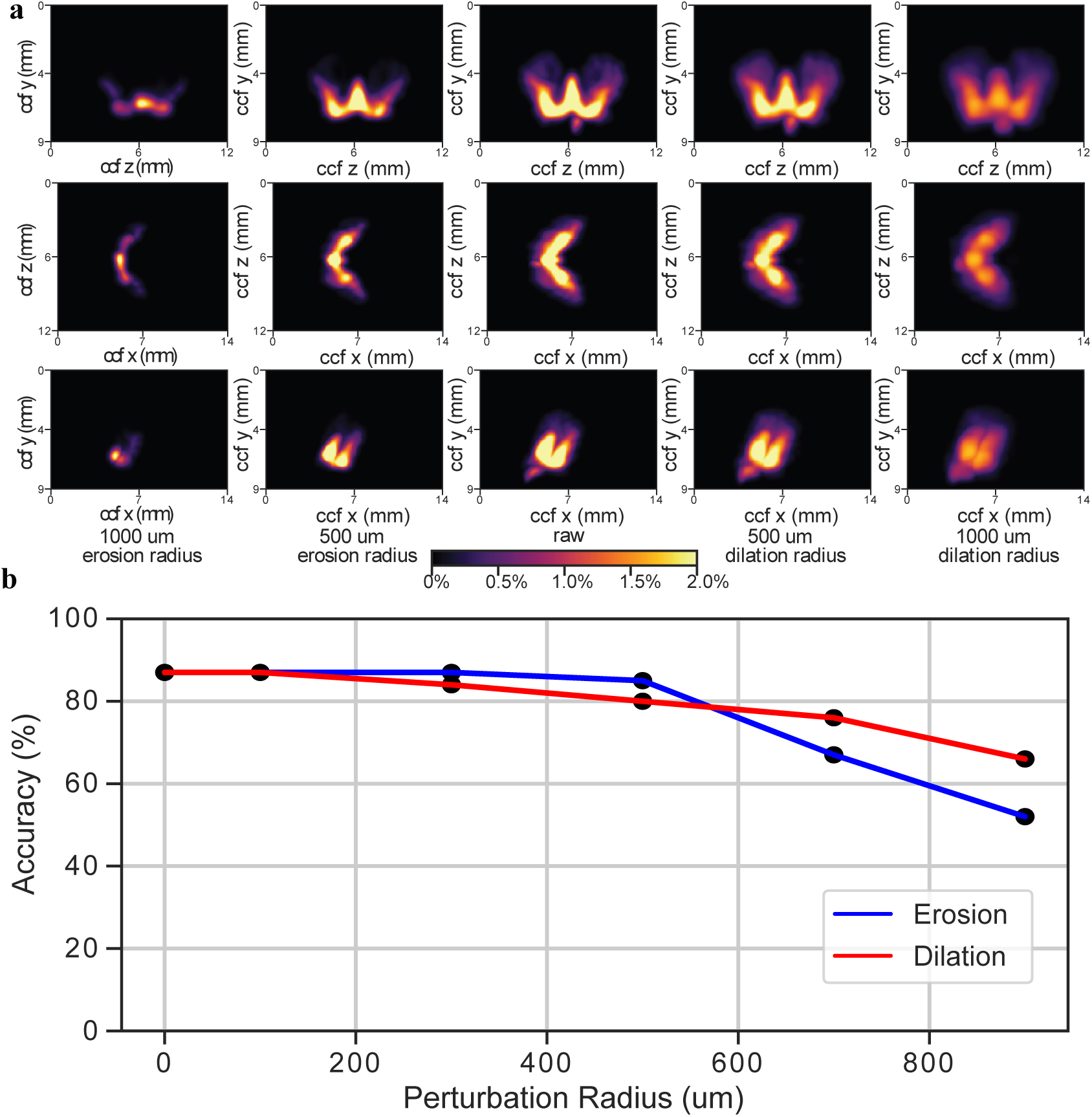
Impact of Priors and Algorithmic Choices on ATLAS/SCALE Performance. **(a)** Representative visualization of the spatial prior for a single cell type, 058 PAL-STR Gaba-Chol. In the central “raw” panel, colors represent the percentage of cells of this type within a given CCF voxel, as determined from reference MERFISH data. The surrounding panels illustrate the effect of applying a 500 µm or 1000 µm erosion radius (left panels) or dilation radius (right panels) to this spatial prior. The color bars indicate the percentage of cells of that type in each voxel. **(b)** Simulation of the sensitivity of ATLAS performance to perturbations in the spatial prior. Classification accuracy is plotted against the perturbation radius for both erosion and dilation. High accuracy is maintained for perturbations up to several hundred micrometers before decreasing relative to performance with the unperturbed “raw” prior.

**Supplementary Figure 4:**
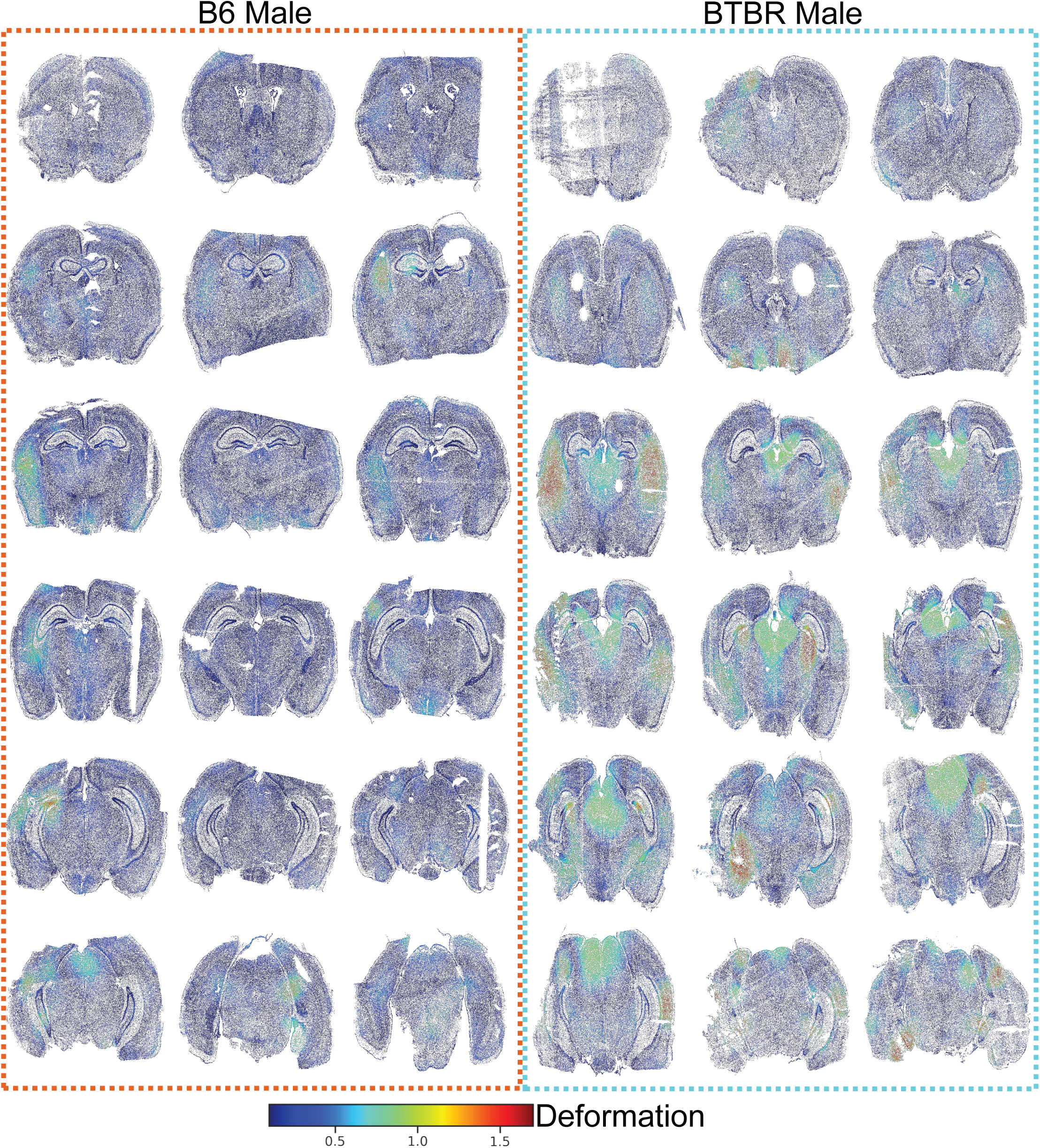
BTBR Morphology Differences. B6 male and BTBR male sections are shown ordered from anterior to posterior. Deformation scores were calculated as the absolute fold change in density for each cell before and after registration to the Common Coordinate Framework. Known morphological changes in BTBR mice, including loss of the corpus callosum and a severely reduced hippocampal commissure, show high deformation scores compared to the relatively low deformation scores in B6 mice.

**Supplementary Figure 5:**
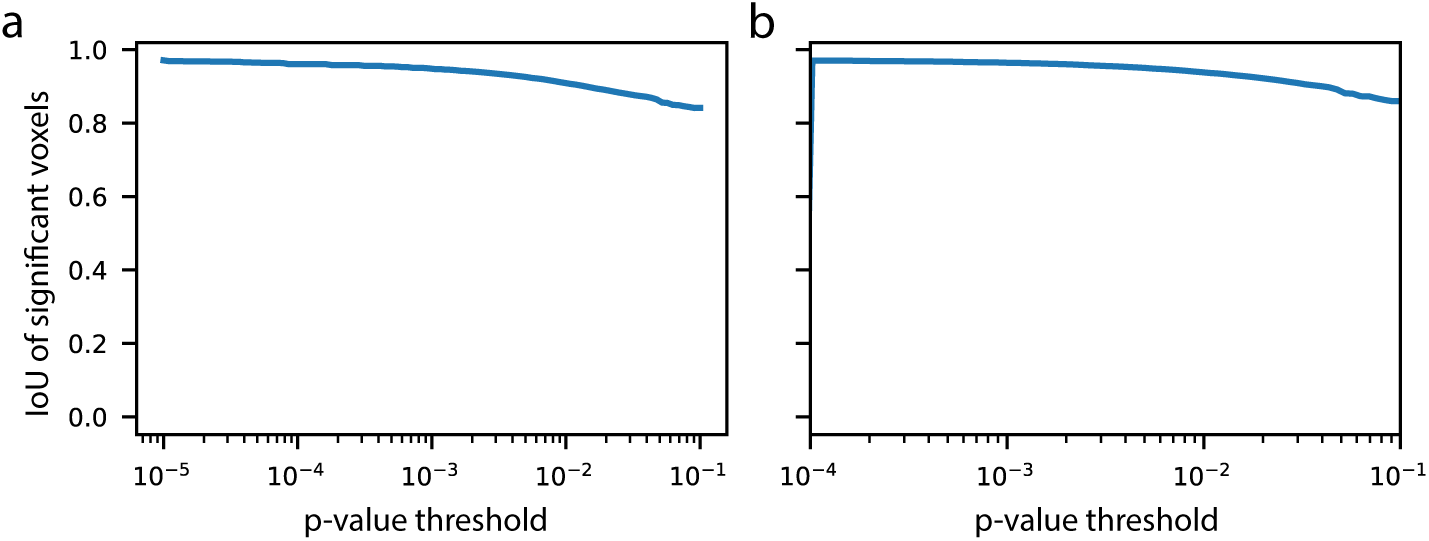
Regional compositional results are largely preserved when SCALE is decoded without spatial priors. Thresholded p-value maps from the Figure 5 regional compositional analysis were compared with maps obtained after repeating SCALE decoding without spatial priors. For each p-value threshold, agreement is shown as the intersection-over-union of significant voxels between the two analyses within the hemisphere shown in Figure 5. (a) B6 male versus BTBR male. (b) B6 male versus B6 female.

**Supplementary Figure 6.**
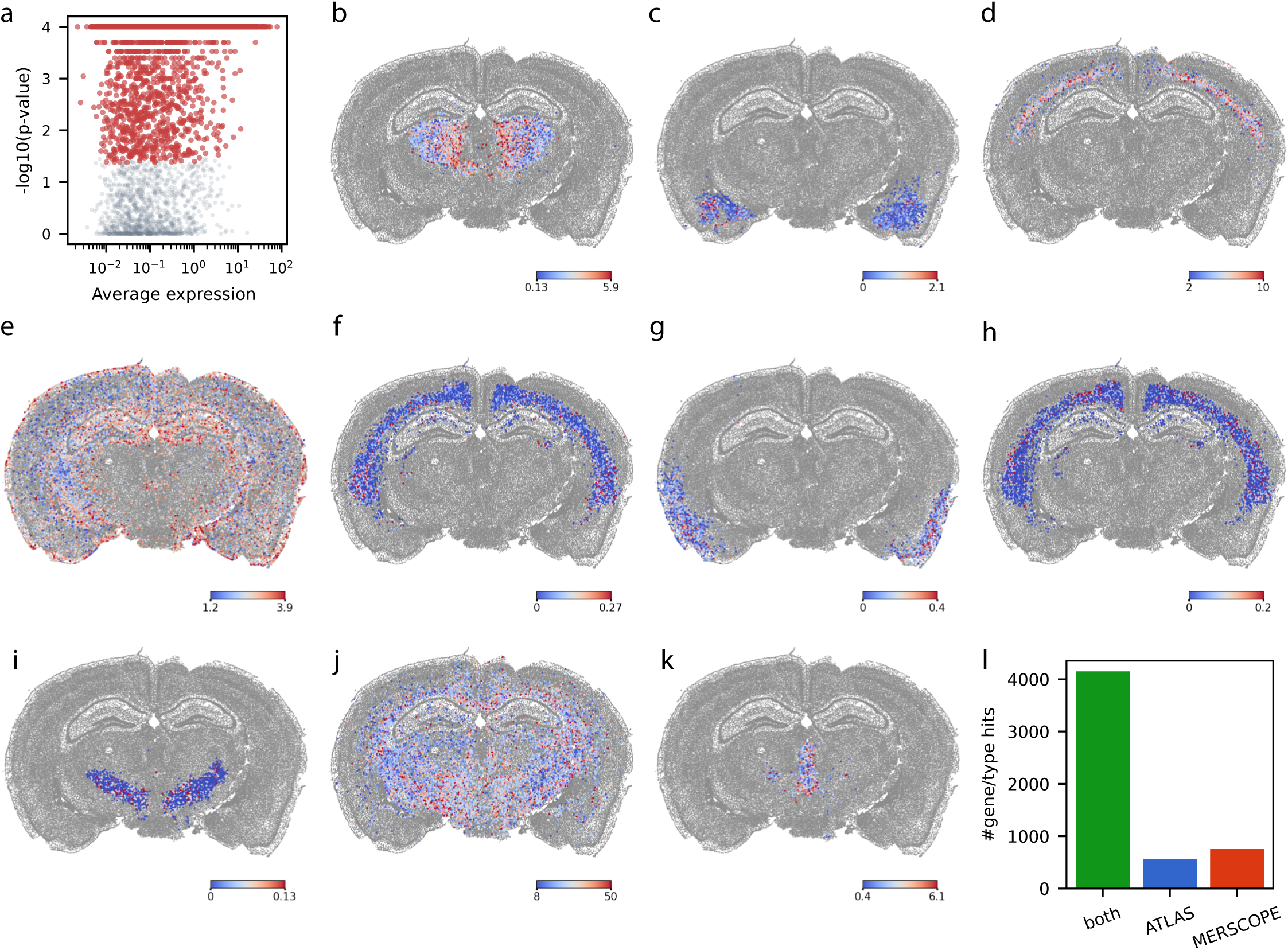
Spatially differential gene expression within cell types. To determine whether ATLAS inference captures spatial variation beyond cell type identity, we tested for spatially differential gene expression within individual cell types using the method of Littman et al. (Mol. Syst. Biol., 2021; see Methods). a. For a single section from a B6 male animal (CCF_X = 7.8), the p-value from the spatial uniformity test is plotted against average gene expression, with each point representing a gene within a given cell type. Red markers show gene/type pairs that differ significantly from a uniform distribution after FDR correction. b–k. Examples of cell-type/gene pairs showing statistically significant non-uniform spatial expression. The panel legend indicates the cell type and gene symbol: b. TH Prkcf Grin2c Glut / Vwc2l c. LA BLA BMA PA Glut / Cpa6. d. L6 CT CTX Glut / Epha4. e. Astro TE / Pou2f2. f. L6 CT CX Glut / Pthr2r. g. L2/3 IT PIR ENTI Glut / Tfap2d. h. L6 CT CTX Glut / Oprk1. i. ZI Pax6 Gaba / Npbwr1 j. Oligo NN / St18. k. SPA SPFm SPFp POL PIL PoT Sp9 Glut / Epha4. I. Comparison of cell-type-specific spDEGs found in the ATLAS section and a matched MERSCOPE section.

**Supplementary Figure 7.**
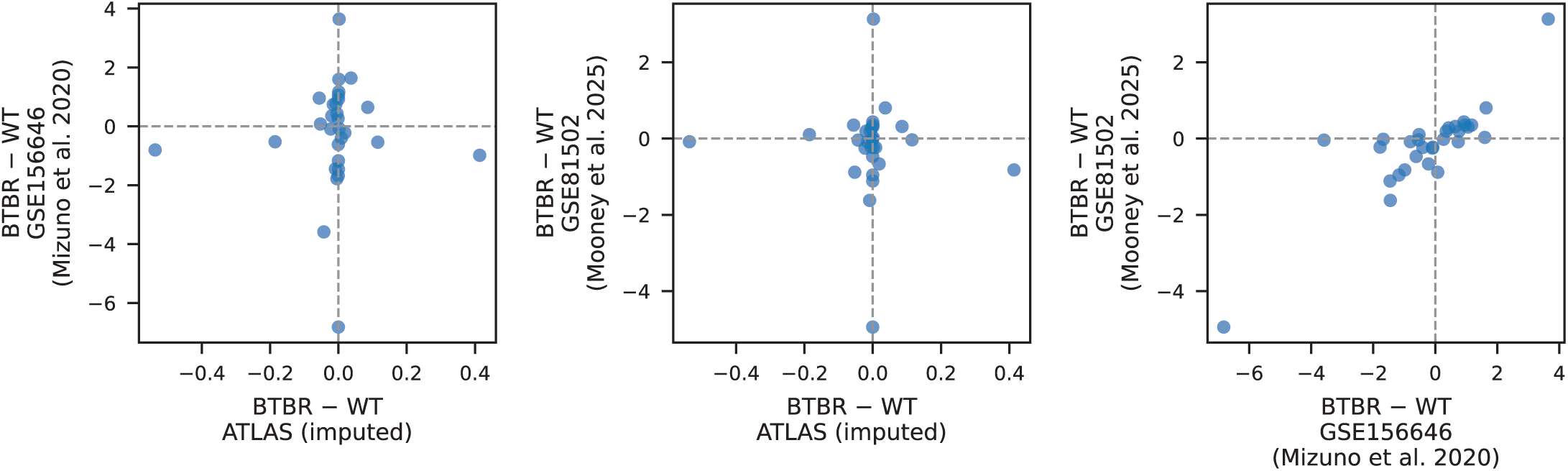
WT-reference ATLAS imputation does not recover BTBR-associated differential expression. Comparison of BTBR versus WT gene-expression differences for 30 genes previously reported as altered in BTBR mice by Mizuno et al. (2020) and Mooney et al. (2025). Each point represents one gene, plotted as the BTBR minus WT expression difference in the indicated datasets. ATLAS-imputed expression changes showed essentially no agreement with direct measurements from either external dataset, with R^2^=0.00 for both comparisons. In contrast, the two direct-measurement datasets were substantially concordant with each other (R^2^=0.72). These results indicate that gene-expression values imputed from a WT reference atlas do not reliably recover BTBR-associated differential expression.

